# Dynamic relocalization of the cytosolic type III secretion system components prevents premature protein secretion at low external pH

**DOI:** 10.1101/869214

**Authors:** Stephan Wimmi, Alexander Balinovic, Hannah Jeckel, Lisa Selinger, Dimitrios Lampaki, Emma Eisemann, Ina Meuskens, Dirk Linke, Knut Drescher, Ulrike Endesfelder, Andreas Diepold

## Abstract

Many bacterial pathogens use a type III secretion system (T3SS) to manipulate host cells. Protein secretion by the T3SS injectisome is activated upon contact to any host cell, and it has been unclear how premature secretion is prevented during infection. We found that in gastrointestinal pathogens, cytosolic injectisome components are temporarily released from the proximal interface of the injectisome at low external pH, preventing protein secretion in acidic environments, such as the stomach. In *Yersinia enterocolitica*, low external pH is detected in the periplasm and leads to a partial dissociation of the inner membrane injectisome component SctD, which in turn causes the dissociation of the cytosolic T3SS components. This effect is reversed upon restoration of neutral pH, allowing a fast activation of the T3SS at the native target regions within the host. These findings indicate that the cytosolic components form an adaptive regulatory interface, which regulates T3SS activity in response to environmental conditions.

## Introduction

In order to proliferate in contact to eukaryotic host cells, both symbiotic and pathogenic bacteria have developed methods to influence host cell behavior. The type III secretion system (T3SS) injectisome is a molecular machinery used by various pathogenic bacterial genera including *Salmonella, Shigella*, pathogenic *Escherichia, Pseudomonas* and *Yersinia* to deliver molecular toxins – effector proteins – directly into the eukaryotic host cells. While the effectors differ among the different bacterial species (Büttner, 2012), and can have different functions in modulating the cytoskeleton, invading and escaping host cells or endosomes, or inducing host cell death (Hueck, 1998; Coburn *et al*, 2007), the structural proteins of the injectisome are highly conserved (Deng *et al*, 2017; Wagner *et al*, 2018). An extracellular needle is formed by helical polymerization of a T3SS-exported protein, and ends in a pentameric tip structure. At the proximal end, the needle is anchored by two multimeric membrane rings that span the outer and inner membrane. Additionally, the inner membrane (IM) ring encloses the export apparatus. At the cytosolic interface of the injectisomes, four soluble T3SS components (SctK/Q/L/N)^a^ interact to form six pod structures (Hu *et al*, 2015, 2017) (Fig. 1A).

**Fig. 1.**
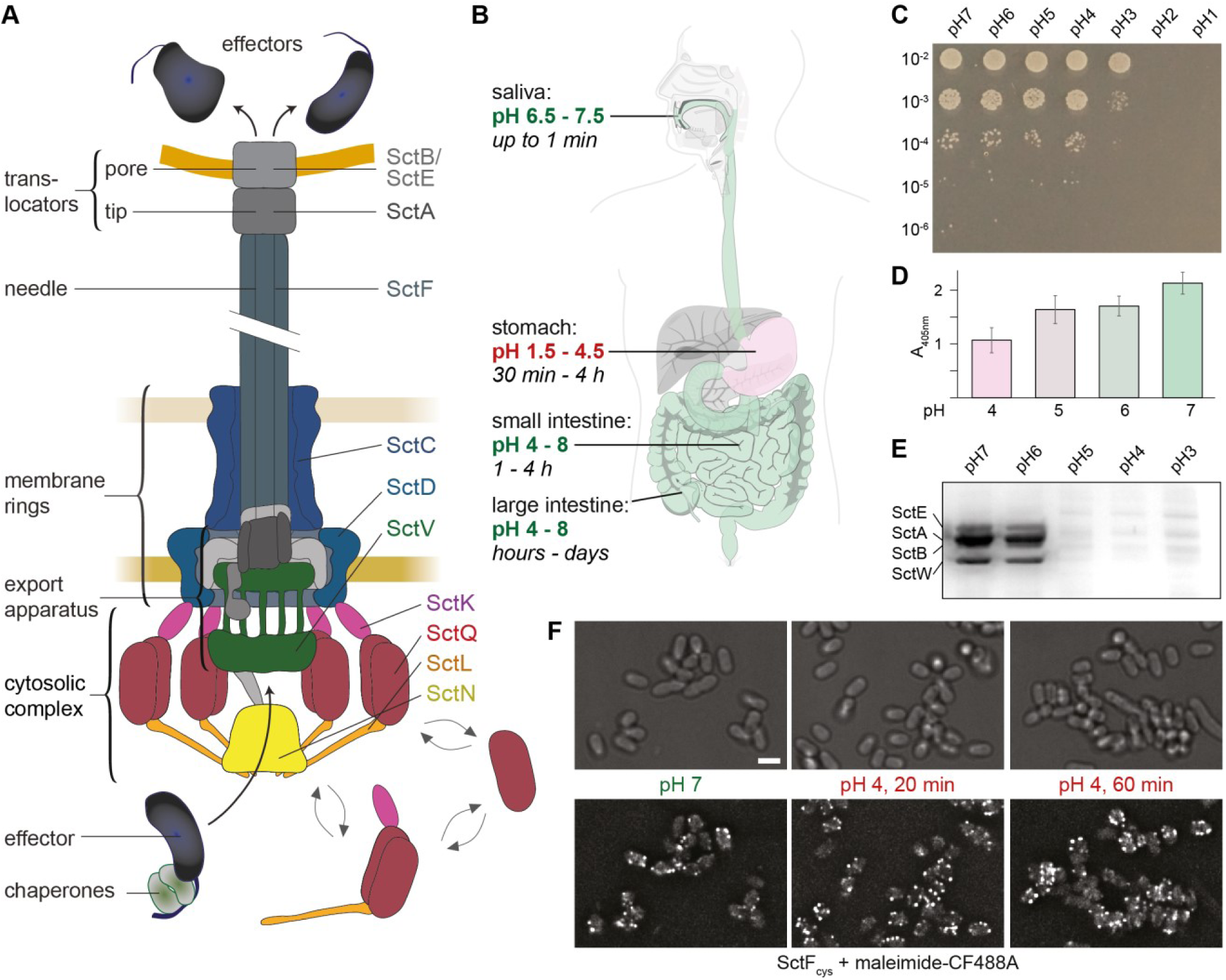
The *Yersinia enterocolitica* T3SS and its function and stability at low pH. (**A**) Schematic representation of the active T3SS injectisome (modified from (Diepold & Wagner, 2014)). Left side, description of main substructures; right side, names of T3SS components studied in this manuscript. Double arrows indicate the exchange of the cytosolic subunits or subcomplexes between the cytosolic and the injectisome-bound state. (**B**) pH ranges and typical retention times at different parts of the gastrointestinal system (Evans *et al*, 1988; McClements & Li, 2010). Digestive tract image based on the public domain template https://commons.wikimedia.org/wiki/File:Digestive_system_diagram_de.svg. (**C**) Dilution drop test of *Y. enterocolitica* cultures incubated at the indicated pH for 15 minutes. (**D**) Binding of the *Y. enterocolitica* adhesin YadA to collagen at the indicated pH values. Absorption at 405 nm resulting from Ni^2+^-HRP binding to YadA-His_6_ incubated with plate-absorbed calf collagen type I. *n* = 3, error bars denote standard deviation. (**E**) *In vitro* secretion assay showing the export of native T3SS substrates (indicated on left side) in strain IML421 at the indicated external pH values. Black and white scan of a coomassie-stained SDS-PAGE gel; supernatant of 3*10^8^ bacteria per lane. Secretion assay of strain containing effectors, see Suppl. Fig. 2A. (**F**) Staining of T3SS needles at the different indicated external pH values; time indicates the duration of pH 4 treatment. A strain expressing the mutated needle subunit SctF_cys_ was covalently labeled with maleimide-CF488A. Scale bar, 2 μm.

The soluble T3SS components SctK, SctQ, SctL and SctN interact in a linear fashion (Jackson & Plano, 2000), and the presence of all four proteins is needed for their assembly at the injectisome, and subsequently for effector secretion (Diepold *et al*, 2010, 2017). SctK/Q/L form a high molecular weight complex that has been shown to bind chaperones and effectors (Lara-Tejero *et al*, 2011). Since the cytosolic components do not co-purify with the rest of the needle, their structural arrangement has only been revealed by recent *in situ* cryo-electron tomography (Hu *et al*, 2015; Nans *et al*, 2015; Makino *et al*, 2016; Hu *et al*, 2017), which showed the formation of six pod structures at the cytosolic interface of the injectisome. The connection between these pod structures and the membrane rings is established by the cytosolic component SctK, which binds to SctD. In the presence of SctK, the cytosolic domains of SctD (SctD_C_) rearrange and in the case of the *Salmonella* SPI-1 T3SS form six discrete patches of four SctD_C_ each that interact with one SctK protein, which in turn connects to SctQ, SctL, and SctN (Hu *et al*, 2017; Tachiyama *et al*, 2019).

In addition to forming the injectisome-bound pod structures, the cytosolic components exist in a freely diffusing cytosolic state, with proteins exchanging between the two states (Diepold *et al*, 2015). In the cytosol, SctK/Q/L/N form a dynamic adaptive network, with a variety of complexes of different stoichiometries that change their composition in response to different external conditions (Diepold *et al*, 2017; Bernal *et al*, 2019). Strikingly, protein interactions and exchange rates amongst the cytosolic components correlate with the activity of the injectisome (Diepold *et al*, 2017; Rocha *et al*, 2018; Diepold *et al*, 2015). However, the precise role of the cytosolic components in the secretion process remains unclear to this date.

*Yersinia enterocolitica* is an extracellular gastrointestinal pathogen that employs its T3SS to downregulate immune responses and prevent inflammation after the penetration of the intestinal epithelium. For initial attachment to the epithelium, the bacteria employ a number of adhesins, with *Yersinia* adhesin A (YadA) as the key factor for establishing an infection (Meuskens *et al*, 2019). YadA interacts with a variety of extracellular matrix molecules, including collagen and fibronectin (Mühlenkamp *et al*, 2015). YadA length is tightly linked to the length of the injection needle of the T3SS, and establishes close contact to the host cells enabling injection (Mota *et al*, 2005). *Y. enterocolitica* is usually taken up with contaminated food or water. The shift from environmental to host temperature (37°C) induces expression and assembly of the injectisomes as well as YadA (Tertti *et al*, 1992; Cornelis, 2006). The bacteria then have to pass the acidic environment of the stomach. During that time, the injectisome can already be present and ready for secretion of translocators (which form a pore in the host membrane and its connection to the needle, Fig. 1A) and effectors. Since the T3SS readily translocates cargo into any host cell type it adheres to, including immune cells, epithelial cells and even red blood cells (Clerc *et al*, 1986; Håkansson *et al*, 1996), a distinct mechanism is needed to prevent premature activation of the T3SS during the passage of the stomach, which would result in a loss of valuable resources, or even elicit immune responses.

We hypothesized that the acidic environment in the stomach (external pH of 1.5 – 4.5 (Evans *et al*, 1988; McClements & Li, 2010), Fig. 1B) may be detected by bacteria to inhibit effector secretion under these conditions. Here, we investigated this hypothesis with a combination of fluorescence microscopy, single particle tracking, and functional assays. Our findings show that although parts of the T3SS, including the extracellular needle, are stable at low pH, a set of cytosolic T3SS components temporarily unbind from the injectisome at low external pH. The reversible dissociation corresponds to a temporary suppression of effector secretion by the T3SS. This mechanism prevents premature activation of the T3SS, while ensuring a quick reactivation of the T3SS, once a pH-neutral environment is reached. Our data provide a striking example for how bacteria apply protein dynamics to adapt the function of a large macromolecular machine essential for virulence to external conditions.

## Results

### Resistance of *Y. enterocolitica* and its T3SS needles to low external pH

To investigate to which degree *Y. enterocolitica* can withstand a drop in external pH, cultures of *Y. enterocolitica* IML421asd (a strain lacking the six main virulence effectors YopH,O,P,E,M,T) (Kudryashev *et al*, 2013) in exponential growth phase were exposed to different pH for 15 minutes at 28°C. A subsequent dilution series on neutral pH agar showed nearly complete survival down to pH 4 (Fig. 1C). Likewise, bacterial cultures remained at a constant optical density at pH 4 at 37°C for at least 90 min, and recovered their growth upon restoration of neutral pH (Suppl. Fig. 1). Together, these results show that *Y. enterocolitica* tolerates temporary incubation down to pH 3, with no or very little fitness decrease at pH 4 and above.

*Y. enterocolitica* adheres to host cells and other surfaces by adhesins, most importantly the trimeric adhesin YadA and invasin (Keller *et al*, 2015; Mühlenkamp *et al*, 2015; Leo *et al*, 2015). We thus tested whether low pH prevents the binding of YadA to collagen and more generally, of *Y. enterocolitica* cells to surfaces. Binding could be established at low pH in both cases (Fig. 1D, Suppl. Videos 1-2), albeit at a reduced level (about 50% binding at pH 4). These results suggest that to avoid protein translocation into non-host cells, secretion itself might be prevented at low pH.

To test under which conditions *Y. enterocolitica* secretes T3SS substrates, we performed an *in vitro* secretion assay where *Y. enterocolitica* cells primed for secretion were subjected to secreting media in the range from pH 8 to pH 4. Indeed, we observed that secretion did not occur at low pH (Fig. 1E, Suppl. Fig. 2A). Lack of secretion is not due to lower protein synthesis at pH 4 (Suppl. Fig. 2B), suggesting a specific mechanism to suppress secretion at low external pH.

But how does *Y. enterocolitica* prevent secretion at low pH? To determine if the absence of secretion is due to a complete disassembly of injectisomes at that pH, we visualized the needles at different pH values and over time by labeling an introduced cysteine residue with a maleimide-linked dye (Milne-Davies *et al*, 2019). The needles were stable at pH 4 over continued time periods (Fig. 1F, Suppl. Fig. 3).

Taken together, *Y. enterocolitica* as well as its injectisome needles can withstand low external pH conditions, but still translocators and effectors are not secreted.

### Association of the dynamic cytosolic T3SS components to the injectisome is temporarily suppressed at low external pH

We have recently found that the cytosolic T3SS components (SctK/Q/L/N) form a dynamic network, where protein exchange is connected to the function of the injectisome (Fig. 1A) (Diepold *et al*, 2015, 2017). Hence, we wondered whether these dynamic components could be involved in the inhibition of secretion at low pH. To investigate this question, we performed flow-cell-based total internal reflection fluorescence (TIRF) microscopy with functional N-terminal fluorescent protein fusions of the cytosolic components, expressed at native levels^b^: EGFP-SctK, EGFP-SctQ, EGFP-SctL and EGFP-SctN (Fig. 1A) (Diepold *et al*, 2010, 2017). At neutral or near-neutral pH, the cytosolic components localized in foci at the bacterial membrane, which represent their injectisome-bound state (Diepold *et al*, 2010). However, at an external pH of 4, all cytosolic components lost this punctuate localization, and the proteins relocated to the cytosol (Fig. 2 ABC). The relocation remained stable over time at low external pH (Suppl. Fig. 4). Strikingly, this phenomenon was reversible: Upon exposure to neutral external pH, the foci recovered within a few minutes (Fig. 2ABC). This effect was observed both under secreting and non-secreting conditions (Suppl. Fig. 5A), and was independent of the fluorophore or visualization tag that was used (Suppl. Fig. 5B). Dissociation and re-association of the cytosolic T3SS components in response to the external pH was reversible for several cycles (Suppl. Video 3). This reversible response to low pH was also observed both under secreting and non-secreting conditions (presence of 5 mM CaCl_2_ and EGTA, respectively) (Suppl. Fig. 6).

**Fig. 2.**
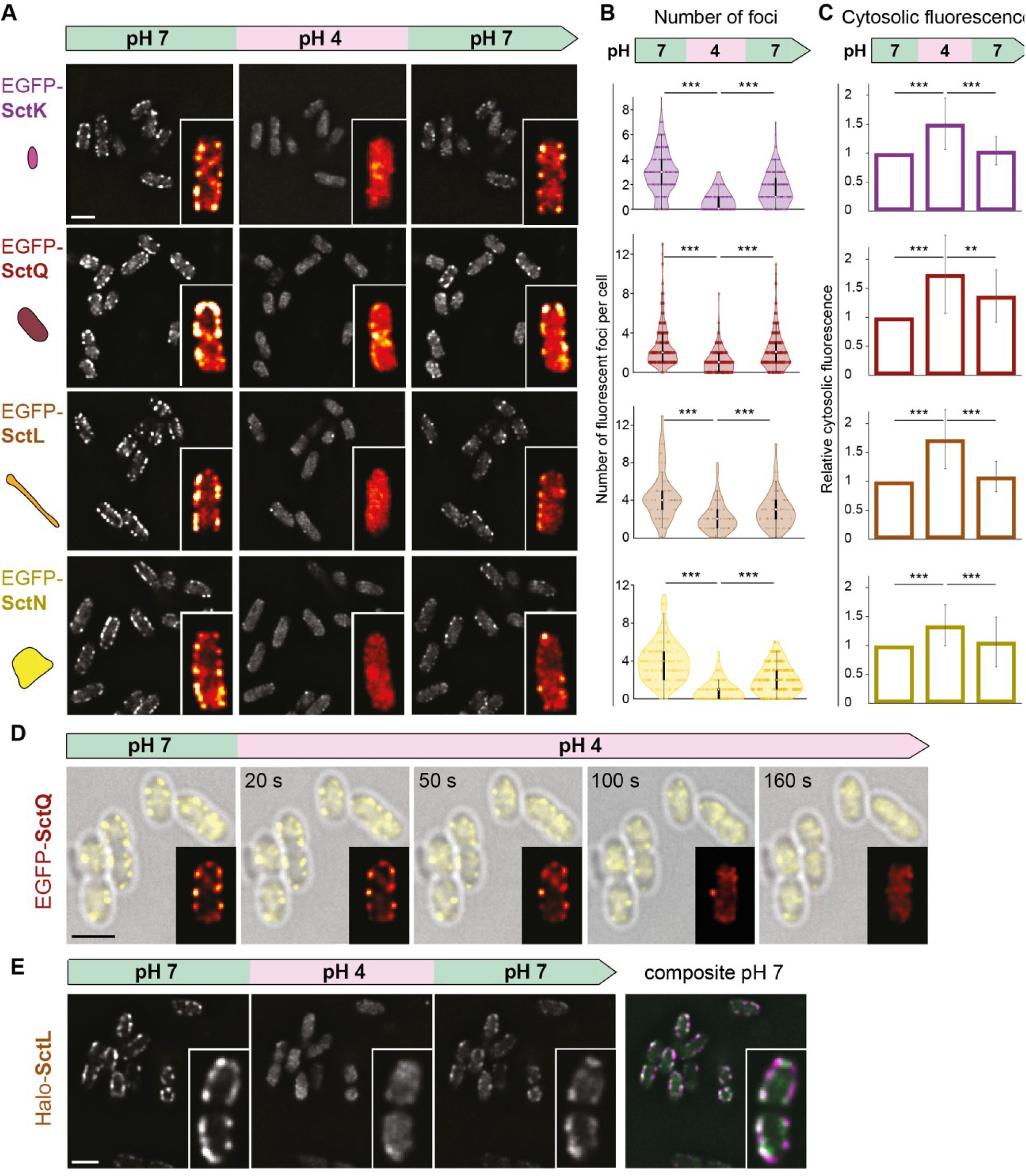
The cytosolic T3SS components temporarily dissociate from the injectisome at low external pH. (**A**) Fluorescent micrographs of the indicated proteins in live *Y. enterocolitica*, consecutively subjected to different external pH in a flow cell. Images were taken under secreting conditions, 10 minutes after bacteria were subjected to the indicated pH. Insets, enlarged single bacteria, visualized with the ImageJ red-hot color scale. (**B**) Quantification of foci per bacterium for the strains and conditions shown in panel (A). *n* = 324, 320, 220, 117 foci (from top to bottom) from 2-3 fields of view from a representative experiment. (**C**) Quantification of mean cytosolic fluorescence for the strains and conditions shown in panel (A). *n* = 50 cells from 2 independent experiments per strain. For (B) and (C), error bars denote standard deviation; */**/***, p<0.05/0.01/0.001 in homoscedastic two-tailed t-tests. The values at pH 7 before and after incubation at pH 4 differ statistically significantly for (B) (***), but not for (C) (p>0.05), except for SctQ (***) and SctL (*). (**D**) Kinetics of EGFP-SctQ dissociation after pH shift from 7 to 4. Overlay of phase contrast (grey) and fluorescence images (yellow); insets, enlarged single bacterium, visualized with the ImageJ red-hot color scale. (**E**) Fluorescence micrographs of Halo-SctL, labeled with JF-646 prior to the first image, during consecutive incubation at different external pH as in (A). Right, overlay of fluorescence distribution at pH prior to and after pH 4 incubation (magenta and green, respectively). Scale bars, 2 μm

To determine the kinetics of association and dissociation of SctQ, we monitored EGFP-SctQ foci in a microscopy flow cell after a pH drop from 7 to 4 and *vice versa*. Foci gradually disappeared within two minutes under secreting condition (Fig. 2D, Suppl. Fig. 7, Suppl. Video 4). Upon restoration of neutral external pH, the first distinct foci were visible after 50 seconds, and further increased in abundance and brightness over time (Suppl. Fig. 8, Suppl. Video 5).

To test whether the recovery of foci was due to the synthesis of new proteins or if the previously bound proteins rebind to the injectisomes, we covalently labeled the pool of Halo-tagged SctL with the Janelia Fluor JF-646 Halo ligand dye (Grimm *et al*, 2017) prior to the incubation at pH 4 to ensure that only the protein pool present at the time of labelling was fluorescent. Similar to the previous experiments, we observed a reversible loss of fluorescent foci at the membrane and an increase in cytosolic signal at pH 4 and rebinding of the fluorescent proteins upon changing pH to 7 (Fig. 2C). Also, an overlay of the micrograph at pH 7 before and after the incubation at pH 4 further indicated that most foci reform at the same position as they previously appeared (Fig. 2E), and colocalize with the T3SS needles (Suppl. Fig. 9).

Taken together, our data indicate that the cytosolic components reversibly dissociate from the injectisome at low external pH. Upon exposure to neutral pH, proteins from the same pool rebind at the cytosolic interface of the injectisome, forming the potential basis for a regulatory mechanism for the prevention of secretion at low pH.

### Molecular mechanism of extracellular pH sensing

What is the molecular basis for the dissociation of the cytosolic T3SS components at low external pH? To find out whether the pH is sensed intracellularly, we first tested the impact of the changed external pH on the cytosolic pH, using a ratiometric pHluorin GFP variant (pHluorin_M153R_ (Miesenböck *et al*, 1998; Morimoto *et al*, 2011)) as a pH sensor. Upon changing the external pH from 7 to 4, the cytosolic pH dropped to a mildly acidic value (pH 6.3-6.4). This cytosolic pH was retained for at least 30 minutes at external pH 4, but quickly recovered upon re-establishment of neutral external pH (Fig. 3A).

**Fig. 3.**
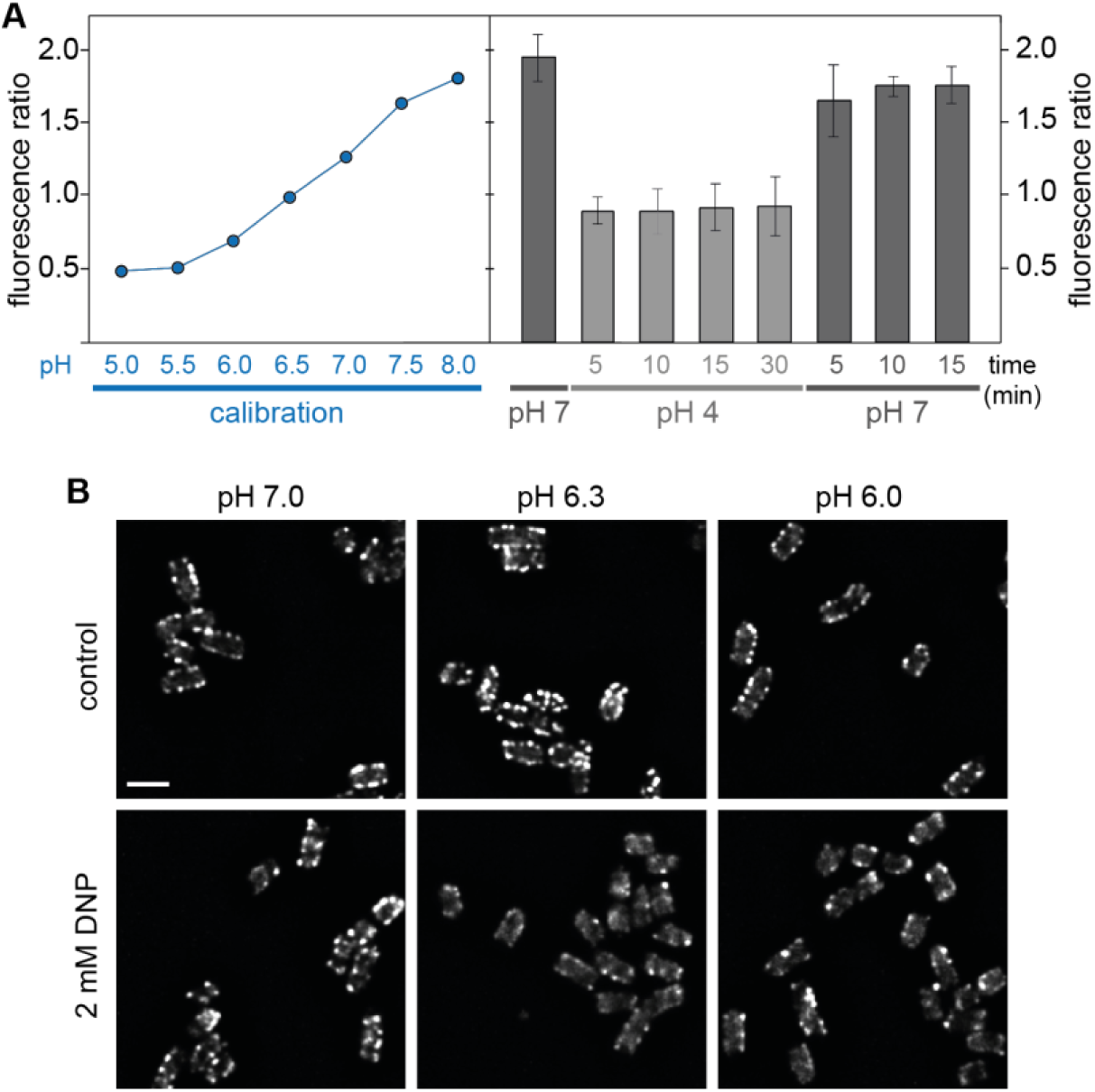
Low external pH leads to a small drop in cytosolic pH, which does not induce the dissociation of cytosolic T3SS components. (**A**) Left, calibration of (Ex_390nm_ / Ex_475nm_) fluorescence ratio of purified pHluorin_M153R_ for the indicated pH values. Technical triplicate, error bars too small to display. Right, determination of cytosolic pH upon changing the external pH from 7 (first column) to 4 (columns 2-5) and back (columns 6-8). Fluorescence ratio (Ex_390nm_ / Ex_475nm_) of bacteria expressing cytosolic pHluorin_M153R_. *n* = 4, error bars denote standard deviation. (**B**) Fluorescence distribution of EGFP-SctQ in live *Y. enterocolitica* at indicated external pH in absence (top) or presence (bottom) of the ionophore 2,4-dinitrophenol (DNP). At pH 4.0, no foci are present. Scale bar, 2 μm.

To test if this mild drop in cytosolic pH directly causes the dissociation of the cytosolic complex, we treated bacteria with the proton ionophore 2,4-dinitrophenol, which attunes the cytosolic pH to the external pH (Hong *et al*, 1979; Dechant *et al*, 2010; Petrovska *et al*, 2014) (Suppl. Fig. 10), and visualized the localization of EGFP-SctQ at different pH values. EGFP-SctQ remained localized in foci representing assembled cytosolic complexes at pH 6.3 and below (Fig. 3B), indicating that the observed disassembly of the cytosolic complex is not caused by acidification of the cytosol.

Based on above results, we searched for a periplasmic pH sensor. An obvious candidate is SctD, a bitopic IM protein connecting the outer membrane (OM) ring to the cytosolic components (Fig. 1A) (Ross & Plano, 2011; Hu *et al*, 2017). Earlier studies showed that lack of SctD, or its inability to bind to SctC, leads to a similar cytosolic location of SctK/L/N/Q as observed at an external pH of 4 (Fig. 2A) (Diepold *et al*, 2010, 2017), and that the cytosolic domains of SctD connect to SctK via specific interactions (with four SctD binding to one SctK in *Salmonella* SPI-1) (Hu *et al*, 2017; Tachiyama *et al*, 2019). This suggests that structural rearrangements of SctD could easily lead to dissociation of SctK and, subsequently, all other cytosolic components.

When we tested the behavior of SctD at low external pH, we found that at pH 4, EGFP-SctD foci in the membrane became less intense than at pH 7, with a concomitant increase in fluorescence throughout the membrane. In contrast to the cytosolic components however, the SctD foci did not completely disappear (Fig. 4A). Importantly, like the cytosolic components, SctD recovered its localization in foci at neutral external pH (Fig. 4A). To study this unique phenotype in more detail, we performed single-molecule tracking of PAmCherry-SctD proteins in photoactivated localization microscopy (PALM). These experiments revealed that at an external pH of 7, more than 90% of the SctD molecules in the IM were static; by contrast, at an external pH of 4, more than 40% of the SctD molecules became mobile within the membrane (Fig. 4B, Suppl. Fig. 11), indicating a dissociation from the core injectisome structure at low external pH. Indeed, the large T3SS export apparatus component SctV-EGFP retained its localization within foci at pH 4; however, these foci (probably representing the SctV nonamers) were mobile within the membrane (Fig. 4C). The same behaviour has been detected for SctV-EGFP in the absence of SctD (Diepold *et al*, 2011), supporting the notion that at an external pH of 4, SctV is released from the SctD structure.

**Fig. 4.**
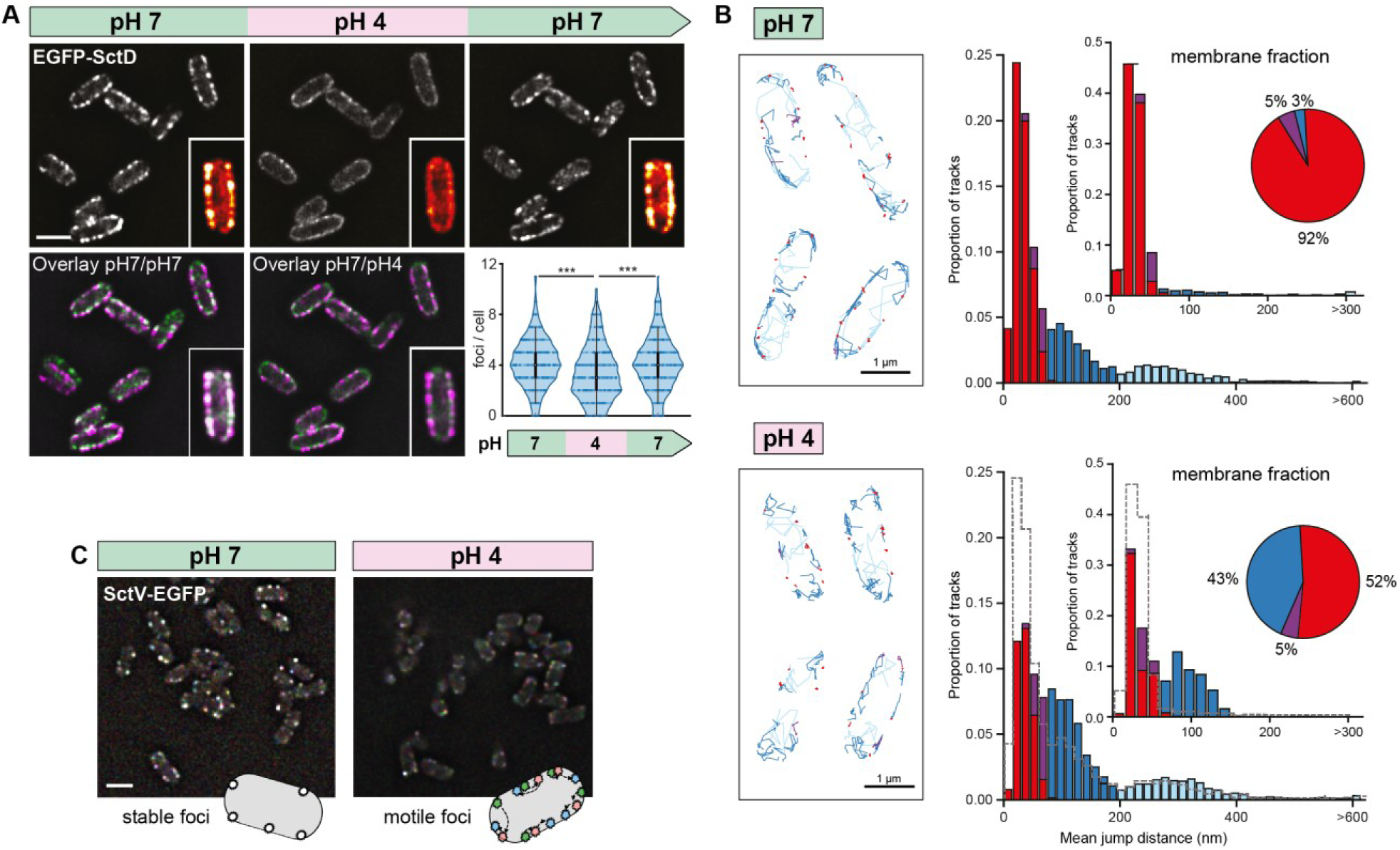
The bitopic IM protein SctD react to low external pH. (**A**) Fluorescent micrographs of EGFP-SctD in live *Y. enterocolitica*, consecutively subjected to different external pH in a flow cell. Images were taken under secreting conditions, 10 minutes after bacteria were subjected to the indicated pH. Insets, enlarged single bacteria, visualized with the ImageJ red-hot color scale. Bottom, overlays of fluorescence at pH 7 before pH change (magenta) and pH 7 after pH change or pH 4 (green). Bottom right, quantification of foci per bacterium. *n* = 1384-1709 foci in 408 cells per condition from a representative experiment. (**B**) PAmCherry-SctD dynamics in exemplary living *Y. enterocolitica* (left) and histograms (right) of the mean jump distances (MJD) of PAmCherry-SctD trajectories weighted by the number of jump distances (# JDs) used for calculating each MJD. Only trajectories with more than 6 one-frame jumps are shown and included into the analysis. Upper panel was measured at pH 7, lower panel at pH 4. Trajectories are assigned into two diffusive states: static (red) and mobile (violet and blue fractions) based on the experimental localization precision. The mobile trajectories are sorted into three MJD categories: lower than 60 nm (violet), lower than 195 nm (dark blue) and higher than 195 nm (light blue). Counts were normalized to the total number of trajectories. At pH 7 we acquired 29,859 trajectories (60% static, 2% mobile below 60 nm MJD, 25% mobile from 60 to 195 nm MJD, 13% mobile faster than 195 nm MJD; 19,473 membrane-bound trajectories). At pH 4 we acquired 33,036 trajectories (34% static, 4% mobile below 60 nm MJD, 44% mobile from 60 to 195 nm MJD, 18% mobile faster than 195 nm MJD; 21,600 membrane-bound trajectories). The bin size of 15 nm was calculated using the Freedman-Diaconis rule. The inset histograms display the statistics of only membrane-bound trajectories. Pie plots display the percentage of each MJD category to the total number of membrane-bound trajectories. The number of trajectories of membrane-bound static PAmCherry-SctD molecules decreases in pH 4 (92% to 52%), while the number of diffusing PAmCherry-SctD with MJDs lower than 195 nm (dark blue bars) increases (3% to 43%). The fast fractions of PAmCherry-SctD molecules of MJDs higher than 195 nm (light blue), which are only visible for the whole cell analysis, remain constant. The grey dashed lines in the pH 4 panels represent the outlines of the pH 7 analysis. (**C**) Spatial stability of SctV-EGFP foci over time at the indicated external pH. Three images of the same focal plane were taken at 10 s intervals. The green channel shows the cell at *t* = 0 s, the blue channel at *t* = 10 s, and the red channel at *t* = 20 s. Scale bars, 2 μm.

### Physiological advantage of temporary suppression of type III secretion at low pH

We reasoned that bacteria could benefit from the dissociation of the cytosolic T3SS components at low external pH to suppress secretion in the stomach, where pH values of 4 and below prevail, which might lead to energy depletion or even elicit immune reactions. If this would indeed be the case, bacteria with a similar infection route, but not bacteria that do not pass through the stomach during normal infection, and therefore are not under evolutionary pressure to suppress T3SS activity at low pH, would be expected to display the same pH dependence for the localization of cytosolic T3SS components. We therefore tested the localization of cytosolic components at low external pH in *Shigella flexneri* and *Pseudomonas aeruginosa. S. flexneri* is a gastrointestinal pathogen that uses an evolutionarily distant T3SS, but like *Y. enterocolitica*, needs to pass the stomach to invade the colonic and rectal epithelium. In contrast, the T3SS of *Y. enterocolitica* and *P. aeruginosa* are closely related (Abby & Rocha, 2012), but the infection strategies of the two species differ. *P. aeruginosa* is not a gastrointestinal pathogen and mainly enters the host body through wounds. In agreement with the hypothesis that the dissociation of cytosolic components at low external pH is a conserved adaptation to the passage of low pH parts of the gastrointestinal system during a normal infection, the fraction of *S. flexneri* GFP-SctN cells (Burgess *et al*, 2020) with foci decreased at pH 4, whereas this was not the case for *P. aeruginosa* EGFP-SctQ cells (Fig. 5, Suppl. Fig. 12).

**Fig. 5.**
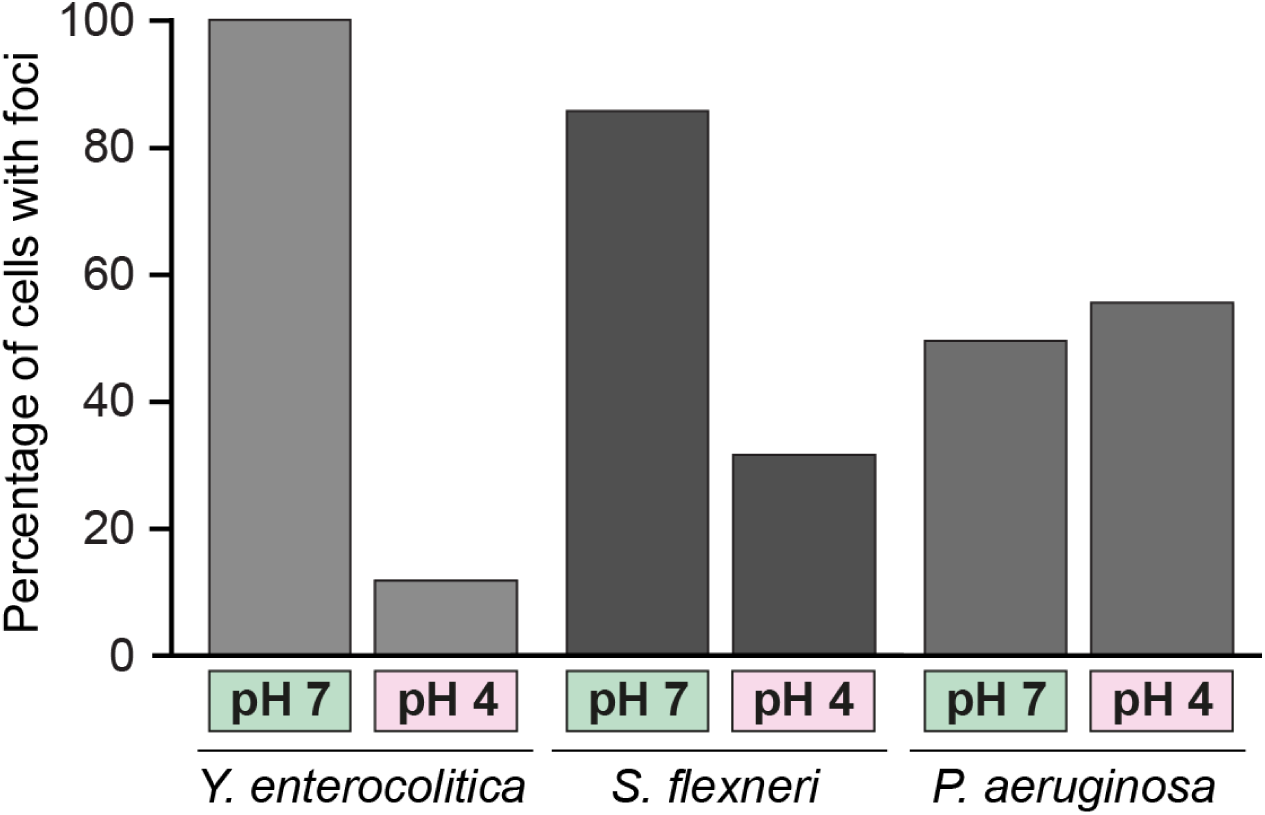
The effect of external pH on the assembly of cytosolic T3SS components is species-specific. Percentage of bacteria with fluorescent foci at the indicated external pH for *Yersinia enterocolitica EGFP-SctQ, Shigella flexneri GFP-SctN*, and *Pseudomonas aeruginosa EGFP-SctQ*, respectively. *n* = 246, 472, 164, 277, 155, 182 bacteria from at least 10 fields of view per condition.

To determine a potential molecular mechanism for this effect, we identified amino acids in the periplasmic domain of SctD that could be protonated at pH 4, but not at pH 7 (Asp, Glu, His), and focused on the subset of these amino acids that differ between *Y. enterocolitica* and *P. aeruginosa* (Suppl. Fig. 13). Showever, single amino acid substitutions, as well as the combination of all four substitutions still led to dissociation of the cytosolic component SctQ, as well as a loss of secretion at pH 4 (Suppl. Fig. 14) suggesting that the effect of the pH might not be conveyed by discrete salt bridges.

After the passage of the stomach, *Y. enterocolitica* arrive in the pH-neutral intestine, where they pass the M cells across the intestinal epithelium. At this point, the injectisome must be ready to manipulate immune cells. To test whether the reversible dissociation of the cytosolic T3SS components supports a fast activation of the T3SS once back at neutral pH, we monitored protein secretion over time after a temporary drop of external pH to 4. We found that secretion was suppressed at low pH, but recovered within the first hour after reaching neutral pH (Fig. 6A). Notably, this recovery is much faster than the onset of effector secretion after *de novo* assembly of the T3SS by a temperature change to 37°C (Fig. 6A), supporting the notion that bacteria benefit from the temporary dissociation of the cytosolic subunits at low pH in two ways: This mechanism suppresses protein secretion at low external pH, while ensuring a fast reactivation upon reaching a pH-neutral environment (Fig. 6B).

**Fig. 6.**
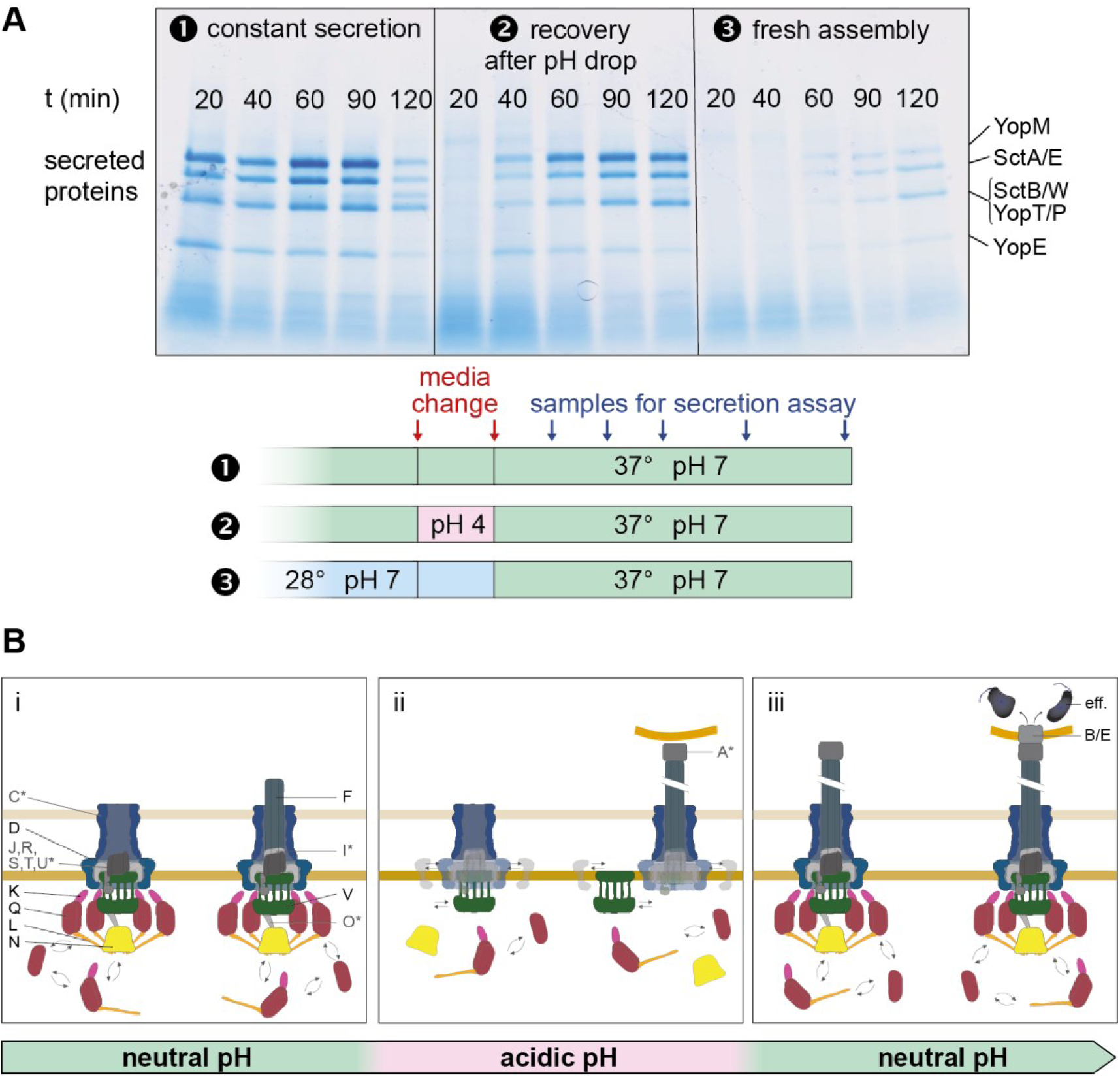
Temporary suppression of T3SS activity at low external pH enables a fast re-activation of secretion. (**A**) *In vitro* secretion assay showing the proteins exported by the T3SS in *Y. enterocolitica* MRS40 at the given time points after the second media change, (1) under constantly secretion-inducing conditions, (2) after a temporary change of the external pH to 4, (3) after incubation at 28°C, where no injectisomes are assembled. Right side, exported effectors, based on (Iriarte & Cornelis, 1998; Diepold *et al*, 2012). (**B**) Model of the pH-dependent suppression of T3SS activity. From left to right: (i) Assembly of the T3SS upon entry into host organisms; cytosolic components bound and exchanging with cytosolic pool. (ii) Prevention of effector translocation upon host cell attachment in the stomach, because cytosolic components are exclusively cytosolic. (iii) Re-association of cytosolic components to the injectisome and effector translocation upon host cell contact in neutral body parts. Letters represent protein identifiers (common Sct nomenclature); eff., effectors. Proteins whose localization was not specifically tested are indicated by asterisks and grey font. Straight double arrows indicate diffusion in membrane; curved double arrows indicate exchange of cytosolic subunits or subcomplexes between the cytosolic and the injectisome-bound state.

## Discussion

On their way through the gastrointestinal system, bacteria encounter a multitude of different pH environments. Importantly, the highly acidic stomach acts as natural barrier for food-borne infections. Gastrointestinal pathogens express factors that facilitate survival in these conditions, such as urease, of which high amounts are exported in *Y. enterocolitica* (Young *et al*, 1996; Hu *et al*, 2009; Heroven & Dersch, 2014; Chen *et al*, 2016; Stingl & De Reuse, 2005). It was not known, however, if and how the activity of the T3SS, an essential virulence factor for many gastrointestinal pathogens, is regulated under these conditions. Although the T3SS target cells are downstream of the stomach for most gastrointestinal pathogens, cells in the gastric mucosa can be accessible and bacteria can attach to host cells at low pH (Fig. 1B, Suppl. Videos 1-2). Our data support the notion that bacteria prevent premature injection into host cells at this stage by directly using the external pH as a cue for the temporary suppression of the T3SS. While parts of the T3SS, including the needle, remain stable, the cytosolic components dissociate from the injectisome in an acidic environment (pH 4 and below). This effect persists at low external pH; however, once the bacteria encounter neutral external pH, both adherence via the adhesin YadA to collagen is significantly increased (Fig. 1D), and the binding of the cytosolic components is restored (Fig. 2). Bacteria conceivably benefit from this mechanism, which prevents premature effector translocation into any eukaryotic cells in contact in the acidic regions of the stomach, an event that would be energetically expensive and might elicit immune responses. Once the pH-neutral intestine is reached, secretion is restarted within 20-40 minutes, which is significantly faster than *de novo* synthesis of injectisomes at this time (Fig. 6).

We found that similarly to *E. coli* (Slonczewski *et al*, 1981; Krulwich *et al*, 2011), *Y. enterocolitica* can partially compensate for acidic external environment, and that at an external pH of 4.0, the cytosolic pH remained at 6.3-6.4 (Fig. 3A). When we used a proton ionophore to create this cytosolic pH at a similar external pH, the cytosolic T3SS components remained bound to the injectisome (Fig. 3B), suggesting that the external pH is sensed outside the bacterial cytosol. A central open question is how exactly the pH change is sensed in the extracytosolic space. A prime candidate to be involved is the bitopic IM component SctD, which could react to a drop in external pH with its periplasmic domain and then transmit this signal to the cytosol. Indeed, our data show that the localization of SctD within the membrane clearly differs between external pH of 7 and 4 (Fig. 4). Interestingly, SctD is one of the least conserved genes in the T3SS, especially in comparison to its direct structural neighbors, the highly conserved SctC secretin ring in the OM, as well as SctJ and the export apparatus proteins in the IM (Diepold & Wagner, 2014). While this low sequence similarity prevented a multi-sequence alignment to identify conserved differences between the T3SS of gastrointestinal and non-gastrointestinal pathogens (except for the closely related *Y. enterocolitica* and *P. aeruginosa*, Suppl. Fig. 12B), it supports the notion that SctD is involved in species-specific adaptation of the T3SS, such as pH sensing. This hypothesis is further substantiated by the finding that in *P. aeruginosa*, which does not pass the stomach during a normal infection, the effect of low pH is significantly restricted, with the majority of EGFP-SctQ foci remaining present at low external pH (Fig. 5). The single or combined mutation of the amino acids in *Y. enterocolitica* SctD that differ from the *P. aeruginosa* homolog and are expected to be protonated at pH 4, but not at pH 7 (histidines 193, 205, 353, 376) to neutral amino acids did not phenocopy the lack of pH sensitivity, suggesting that a broader reversible conformational change at low pH is responsible for the observed effect. This could include the disruption of intermolecular interaction interfaces within or between SctD and the secretin ring SctC. Due to the periplasmic location of any possible fusion protein, SctC could not be directly visualized in this study; however, given the high stability of secretin rings (Rappl *et al*, 2003; Burghout *et al*, 2004; Majewski *et al*, 2018) the interfaces between SctC and SctD, or between neighboring SctD molecules are prime candidates for a pH-dependent temporary dissociation.

Both possible types of pH sensing, direct structural change and rearrangement of intermolecular interactions, are used by biological systems: The *E. coli* membrane-integrated transcriptional regulator CadC senses acidic pH through direct protonation of a charged surface patch in its C-terminal periplasmic domain, and transduces the signal to the cytosol by its N-terminal cytosolic domain (Haneburger *et al*, 2011). Similarly, the *P. syringae* T3SS effector AvrPto transitions from a largely unfolded state in the mildly acidic bacterial cytosol to a well-defined fold in the neutral host cytosol (Dawson *et al*, 2009). Perhaps most strikingly, the highly ordered multimeric structures of bacteriophages undergo large-scale structural changes at low pH, that are reversible (Helenius *et al*, 1980; Mauracher *et al*, 1991; Taylor *et al*, 2002). Indeed, our data indicate a reversible partial delocalization of SctD at low external pH (Fig. 4). The presence of SctD is required for binding of any cytosolic T3SS component (Diepold *et al*, 2010, 2017), most likely through a direct contact of four SctD to one SctK at the cytosolic interface of the IM (Hu *et al*, 2017), indicating that this partial displacement of SctD (Fig. 4) is causal for the dissociation of the cytosolic T3SS components.

The dissociation kinetics of the cytosolic components of the T3SS revealed a dissociation half-time of about one to two minutes under secreting conditions (Fig. 2D, Suppl. Video 3). This value is strikingly similar to the exchange rate of SctQ at the injectisome (Diepold *et al*, 2015), suggesting that at low external pH, primarily the re-association of the cytosolic components is prevented. This adaptation of T3SS function may explain the so-far enigmatic benefit of the observed dynamics of the cytosolic parts of the injectisome during its function. Notably, the observed dissociation of the cytosolic proteins in response to low external pH may not reveal the complete mechanism for suppression of T3SS activity at this pH. The observation that the dissociation of the cytosolic components occurs at a slightly lower pH than the loss of effector secretion, and that the re-initiation of secretion occurs later than the initial recovery of foci indicates that other factors might participate in this phenotype. The slightly delayed activation of secretion after restoration of neutral pH could also be explained by the time required for re-associating SctV to the machinery (the cytosolic components can already bind to the membrane rings in the absence of SctV (Diepold *et al*, 2010)). This delay may, in fact, be beneficial for *Y. enterocolitica*, as it delays the antiphagocytic effects of the T3SS effectors, which otherwise may hamper the passage of *Y. enterocolitica* through the M cells that is required to access the lymphoid follicles of the Peyer’s patches (Cornelis, 2002).

Like the *Y. enterocolitica* T3SS, the intracellular *Salmonella enterica* SPI-2 T3SS is strongly influenced by the external pH; however, the mechanism described for SPI-2 differs from the one described in this study. Injectisome assembly and secretion of the translocon components in SPI-2 are activated by the low pH of the surrounding vacuole (around pH 5.0) (Rappl *et al*, 2003; Yu *et al*, 2010). After this step, neutral pH, most likely indicative of a successfully established connection to the neutral host cytosol, leads to the disassembly of a gatekeeper complex, which in turn licenses the translocation of effectors (Yu *et al*, 2010). Strikingly, a single amino acid exchange in the export apparatus protein SctV governs this pH-dependent effect (Yu *et al*, 2018). Both the sensory and the functional connection between the pH and T3SS activity differ between *Salmonella* SPI-2 and *Y. enterocolitica*, which highlights the high degree of functional adaptability of the T3SS.

In this study, we found that dynamic exchange of cytosolic components of the T3SS enables to suppress the premature activation of the T3SS at low external pH in *Y. enterocolitica*. Strikingly, this effect is conserved in *S. flexneri* and very likely in other gastrointestinal pathogens, but not in *P. aeruginosa*, which does not pass low pH regions during normal infections. Our data highlight that not only the effector repertoire, but also dynamics and function of T3SS are adapted to the specific environment of the respective bacteria. This newly described pathway is a striking showcase for the variety and sophistication of mechanisms that allow bacteria to use different aspects of T3SS assembly and function, including protein dynamics, to tailor the activity of this essential virulence mechanism to their specific needs during infection.

## Material and Methods

### Bacterial strain generation and genetic constructs

A list of strains and plasmids used in this study can be found in Supplementary Table 1. The *Y. enterocolitica* strains used in this study are based on the *Y. enterocolitica* wild-type strain MRS40 (Sory *et al*, 1995), and the strain IML421asd (ΔHOPEMTasd), in which all major virulence effector proteins (YopH,O,P,E,M,T) are deleted (Kudryashev *et al*, 2013). Furthermore, this strain harbors a deletion of the aspartate-beta-semialdehyde dehydrogenase gene which render the strain auxotrophic for diaminopimelic acid (DAP).

Unless stated otherwise, all fusion proteins used were expressed as endogenous fusions from their native location on the pYV virulence plasmid, which were introduced by two step homologous recombination (Kaniga *et al*, 1991).

For cloning and conjugation the *E. coli* strains Top10 and SM10 λpir were used, respectively. All constructs were confirmed by sequencing (Eurofins Scientific). Constructs with single amino acid substitutions in SctD or SctF were created by overlapping PCR using Phusion polymerase (New England Biolabs), and expressed from an arabinose controlled expression vector (pBAD).

### Bacterial cultivation, in vitro secretion assays and fluorescence microscopy

*Y. enterocolitica* day cultures were inoculated from stationary overnight cultures to an OD_600_ of 0.15 and 0.12 for secreting and non-secreting conditions, respectively, in BHI medium (Suppl. Table 2) supplemented with nalidixic acid (35 mg/ml), diaminopimelic acid (80 mg/ml), glycerol (0.4%) and MgCl_2_ (20 mM). Where required, ampicillin was added (0.2 mg/ml) to select for pBAD-based plasmids. For secreting conditions, cultures were additionally supplemented with 5 mM EGTA; for non-secreting conditions, cultures were additionally supplemented with 5 mM CaCl_2_ and filtered through a 0.45 μm filter. Cultures were incubated at 28°C for 90 minutes. At this time point, expression of the *yop* regulon was induced by a rapid temperature shift to 37°C in a water bath and expression of proteins *in trans* was induced by addition of 0.2-1.0% arabinose, as indicated.

For effector visualization or total cell analysis, bacteria were incubated at 37°C between 1-3 hours, as indicated. 2 ml of the culture were collected at 21,000 *g* for 10 min. The supernatant was separated from the total cell fraction and precipitated with trichloroacetic acid (TCA) for 1-8 h at 4°C; proteins were collected by centrifugation for 15-20 min at 21,000 *g* and 4°C, washed once with ice cold acetone and then resuspended and normalized in SDS-PAGE loading buffer. Total cell samples were normalized to 2.5×10^8^ bacteria and supernatant samples to the equivalent of 3×10^8^ bacteria per SDS-PAGE gel lane. Samples were incubated for 5 minutes at 99°C, separated on 12-20% SDS-PAGE gels, and stained with Instant blue (Expedeon).

For fluorescence microscopy, bacteria were treated as described above. After 2-3 h at 37°C, 400 µl of bacterial culture were collected by centrifugation (2,400 *g*, 2 min) and resuspended in 200µl microscopy minimal medium (Suppl. Table 2). Cells were spotted on 1.5% low melting agarose pads (Sigma) in microscopy minimal medium in glass depression slides. Where required, 80 mg/ml DAP for ΔHOPEMTasd-based strains, 0.2-1.0% L-arabinose for induction of *in trans* expression, 5 mM Ca^2+^ for non-secreting conditions, or 5 mM EGTA for secreting conditions were added.

*P. aeruginosa o*vernight cultures were grown at 28°C. For secreting cultures, LB broth (Suppl. Table 2) was supplemented with 20 mM MgCl_2_, 200mM NaCl and 5 mM EGTA and inoculated to an OD_600_ of 0.15 and incubated for 2 h at 37°C. This culture was then used to inoculate a fresh culture containing the same supplements to an OD_600_ of 0.15. Microscopy experiments were then performed 2 h later on a minimal media agar pad and in LB medium.

*S. flexneri* over night cultures where inoculated from a fresh LB-agar plate into BHI broth. On the next day, fresh BHI cultures were inoculated to an OD of 0.12 and grown to an OD of around 1 at 37°C (Burgess *et al*, 2020). 1 ml of bacterial culture was collected by centrifugation (2,400 *g*, 2 min) and resuspended in 100µl microscopy minimal medium (Suppl. Table 2) at the indicated pH. 2 µl of the resuspension were spotted on a minimal medium agar pad adjusted to the indicated pH and then visualized.

### Wide field fluorescence microscopy

Microscopy was performed on a Deltavision Elite Optical Sectioning Microscope (Applied Precision), equipped with a UApo N 100x/1.49 oil TIRF UIS2 objective (Olympus) and a UPlanSApo 100x/1.40 oil objective (Olympus), using an Evolve EMCCD Camera (Photometrics). Samples were illuminated for 0.1 s with a 488 nm laser with a TIRF depth setting of 3440 (TIRF) or for 0.2 s with green LED light (standard oil objective). Excitation and emission filter sets used were 475/28 and 525/48 nm (green), 575/25 and 625/45 nm (red), or 632/22 and 679/34 nm (Cy5). The micrographs where deconvolved using softWoRx 7.0.0 (standard “conservative” settings). Images were then further processed with ImageJ-Fiji. Where necessary, drift correction was performed with the StackReg Plugin. For figures, representative fields of view were selected, and brightness and contrast of the micrographs was adjusted identically within compared images.

Samples sizes and number of replicates for wide field microscopy and all other experiments were determined prior to the experiments, based on experimental feasibility and their potential to draw clear conclusions. These numbers were not changed based on the experimental outcome.

### Flow cell based TIRF microscopy

A microscopy flow cell was manufactured based on (Berg & Block, 1984). Formation of injectisomes was induced under non-secreting conditions. After formation of the injectisomes, bacteria were collected and resuspended in approximately 0.5 volumes of microscopy medium supplemented with 80 mg/ml DAP and 5mM EGTA. The flow cell was pre-incubated with microscopy medium and a bottom coverslip (25 mm No. 1.5; VWR) was attached to the cell. Bacteria were spotted on the coverslip and incubated for 1 min to allow attachment to the glass surface. The cover slip was covered with 60 µl of minimal medium and sealed from the top with an additional cover slip. The flow cell was mounted on the microscopy stage and gravity-driven buffer flow in the chamber was induced using a 1 ml syringe. To exchange the buffer in the flow cell, the tube was quickly relocated to the new buffer. Complete buffer exchange in the flow cell was determined to take 39 ± 5 s (*n* = 7).

### Detection and quantification of fluorescent foci

Detection and quantification of foci was performed by first segmenting the images using the software BiofilmQ. After segmentation, we fluorescent foci were detected within BiofilmQ. The foci were filtered by their intensity above the background intensity of the cell, calculated by subtracting a blurred image.

### Maleimide based needle staining

To visualize the injectisome needles, expression of SctF_S5C_ (Milne-Davies *et al*, 2019) was induced from plasmid. Protein expression was induced with 0.4-1.0% L-arabinose at the temperature shift. After 2-3 h, bacteria were collected and resuspended in 0.2 volumes of microscopy medium supplemented with 5 µM of a CF™488A/633 maleimide dye (Sigma-Aldrich, USA) for 5 min in 100 µl of minimal medium at 37°C. After staining, cells were washed once with 500 µl of minimal medium and spotted on 1.5% agarose pads in the same medium.

### Halo staining with Janelia fluorescent dyes

To stain and visualize a defined pool of T3SS proteins, Halo-labeled proteins were visualized using Janelia Fluor 549-NHS ester and Janelia Fluor 646-NHS ester. 500 µl of bacterial culture were collected by centrifugation (2,400 *g*, 2 min) and resuspended in 100 µl microscopy medium. Bacteria were stained with 0.2 µM Janelia Fluor JF-646/JF-549 dyes at 37°C for 30 min in a tabletop shaker. Afterwards, bacteria were washed twice in 500 µl minimal medium and then visulaized in a flow cell or on 1.5% agarose pads in the same medium.

### Equilibration of cytosolic and external pH

2,4-dinitrophenol (DNP) was diluted in methanol and used at a final concentration of 2 mM in microscopy minimal medium. Cells were grown as described above, and resuspended in DNP containing medium immediately before being spotted on 1.5% agarose pads in the same medium containing DNP.

### Bacterial survival test

Wild-type MRS40 *Y. enterocolitica* were inoculated to an OD_600_ of 0.12 in non-secreting conditions and incubated for 1.5 h at 28°C to reach exponential growth phase. Cells were collected by centrifugation at 2,400 *g* for 2 min. The supernatant was discarded and the collected bacteria were resuspended in a range of pH-adjusted media buffered with 50 mM glycine, 50 mM HEPES and 50 mM MES. These cultures were incubated for 15 min at 28°C and afterwards a dilution series in neutral media was performed. For the visualization, 4 µl of bacterial suspension were spotted on a neutral LB agar supplemented with nalidixic acid and incubated at 28°C overnight.

### Growth curve assays

Wild-type *Y. enterocolitica* MRS40 were inoculated to an OD_600_ of 0.12 in secreting conditions and incubated for 90 min at 28°C. At this point, cells where collected by centrifugation (2,400 *g*, 2 min), and resuspended in 37°C fresh pre-warmed secreting medium buffered with 50 mM glycine, 50 mM HEPES and adjusted to the indicated pH. After 90 min, one batch of cells incubated at pH 4 were collected again and resuspended in fresh buffered secretion medium at pH 7. All strains were incubated for further 90 min. During the experiment, the OD_600_ was measured every 30 min in technical duplicates.

### pHluorin purification and calibration

The protocol described by Nakamura et al. (Nakamura *et al*, 2009) was adapted for bench top purification. Expression of ratiometric GST-pHluorin_M153R_ in *E. coli* DH5αZI was induced in a 500 ml culture with 1 mM IPTG, followed by 4 h incubation at 28°C. Cells were harvested and stored for further processing at −80°C. Next, cells were thawed on ice, resuspended in lysis buffer and disrupted by two passages through a French Press. The lysate was centrifuged (25,000 rpm, 60 min, 4°C) and the supernatant was gently mixed with the previously equilibrated glutathione agarose at 4°C for 1.5 h in a small spinning wheel. The agarose was collected by centrifugation (500 *g*, 5 min) and washed with PBS three times. A thrombin digest was performed in 2 ml PBS while incubating for 60 min on a roll mill. Agarose was removed by centrifugation (500 *g*, 5 min) and the protein was stored at −80°C for further use.

The calibration was performed on a Deltavision Elite microscope. 5 µl of purified pHluorin protein were spotted on a KOH-cleaned microscopy slide in an enclosed compartment (Thermo Fisher Scientific GeneFrame, AB-0577) and incubated for 5 min to ensure attachment to the glass surface. Then, 10 µl of pH-adjusted PBS buffered with 200 mM glycine, 200 mM HEPES, and 200 mM MES were added. Ratiometric pHluorin fluorescence was determined using the DAPI excitation filter (390/18nm) or GFP excitation filter (475/28nm) at 32% illumination intensity, in combination with a GFP emission filter set (525/48nm) with 0.3 s exposure time. The overall fluorescence of the images before deconvolution was determined and corrected for background fluorescence levels, which were measured with an empty slide filled with PBS. The ratio of DAPI/Green vs Green/Green fluorescence was determined for a pH range from pH 8 – pH 5 in 0.5 pH steps.

### Single particle tracking photoactivated localization microscopy (sptPALM)

Bacteria were cultivated under non-secreting conditions as described above. After 2.5 h of incubation at 37°C, the media was changed to pre-warmed (37°C) minimal microscopy medium containing the same supplements as the BHI before. Cells were incubated for 30 min and then washed four times in 5 volumes of pre-warmed (37°C) EZ medium (Supplementary Table 2) supplemented with DAP and 5 mM CaCl_2_. Bacteria were concentrated 2x after the last wash and spotted on a 1.5% agarose pad in EZ medium supplemented with DAP and 5 mM CaCl_2_ on a KOH-cleaned microscopy slide in an enclosed compartment (Thermo Fisher Scientific GeneFrame, AB-0577). Imaging was performed on a custom build setup based on an automated Nikon Ti Eclipse microscope equipped with appropriate dichroic and filters (ET DAPI/FITC/Cy3 dichroic, ZT405/488/561rpc rejection filter, ET610/75 bandpass, Chroma), and a CFI Apo TIRF 100x oil objective (NA 1.49, Nikon). All lasers (405 nm OBIS, 561 nm OBIS; all Coherent Inc. USA) were modulated via an acousto-optical tunable filter (AOTF) (Gooch and Housego, USA). Fluorescence was detected by an EMCCD camera (iXON Ultra 888, Andor, UK) in frame transfer mode and read-out parameter settings of EM-gain 300, pre-amp gain 2 and 30 MHz read-out speed. The z-focus was controlled using a commercial perfect focus system (Nikon). Acquisitions were controlled by a customized version of Micro-Manager (Edelstein *et al*, 2010). Live cell sptPALM experiments were performed on a customized heating stage at 25°C. Live *Y. enterocolitica* PAmCherry-SctD cells were imaged in HILO illumination mode (Tokunaga *et al*, 2008). Applied laser intensities measured after objective were 35 W/cm^2^ (405 nm) and 800 W/cm^2^ (561 nm). Prior to recording each new region of interest (ROI), a pre-bleaching step of 561 nm illumination was applied for 30 seconds to reduce autofluorescence. Videos were then recorded for 2000 frames pulsing the 405 nm laser every 20^th^ imaging frame at 5 Hz with an exposure time of 200 ms per frame. After sptPALM imaging, a bright light snapshot of all illuminated regions was recorded to obtain the bacterial cell shapes.

Single molecule localizations were obtained using *rapidSTORM* 3.3.1 (Wolter *et al*, 2012) and single cells were manually segmented in *Fiji (ImageJ 1*.*51f)* (Schindelin *et al*, 2012). sptPALM data was tracked, visualized and filtered using a customized tracking software written in C++ (***swift***, unpublished software, RG Endesfelder). Trajectories were allowed to have a maximum of 5 frames of gap time (e.g. caused by fluorophore blinking). Trajectories were assigned to their diffusive states (static and mobile) on the basis of the experimental localization precision of about 25 nm (determined by the NeNA method (Endesfelder *et al*, 2014). For trajectories with more than 6 one-frame jumps, the mean jump distance (MJD) was calculated (jumps spanning several frames due to dark times were not used in MJD calculations). The obtained MJDs were weighted by the number of jumps and displayed in a histogram using OriginPro 9.4 (Origin LAB Corporation, USA).

### Y. enterocolitica YadA and cellular adhesion assays

To measure YadA adhesion to collagen at different pH values, an ELISA-like binding assay was performed (Leo *et al*, 2008, 2010; Saragliadis & Linke, 2019). Clear 96 well plates (Sarstedt, Ref. 82.1581) were coated with 100 µl of calf collagen type I (10 µg/ml, ThermoFisher A1064401) in H_2_O with 0.01 M acetic acid for 1h at room temperature (RT). The supernatant was discarded and the wells were blocked with 1% BSA in PBS and afterwards washed three times with 0.1% BSA in PBS. Afterwards, 100 µL of purified YadA head domains with a C-terminal His_6_-tag at a concentration of 10 µg/mL were incubated in the wells. In order to test binding at different pH, the YadA heads were diluted in acetic acid/sodium acetate at pH 4.0 or pH 5.0 and for higher pH values in PBS at pH 6.0 and 7.0. Binding was allowed for 1 h at RT. Afterwards, the plate was emptied and washed three times with 0.1% BSA in PBS. The wells were blocked with 1.0% BSA in PBS for 1h at RT. For detection, Ni-HRP conjugates (HisProbe-HRP, Thermo Fisher Scientific, Ref. 15165) were diluted in 0.1% BSA in PBS and 100 µl per well were incubated for 1 h at RT. The Ni-HRP conjugate solution was discarded and the wells washed three times with PBS. Detection was performed using ABTS substrate (Thermo Scientific. Ref. 34026). Development was allowed for 30 min. Absorption was measured at 405 nm on a plate reader (Biotek Synergy H1).

Glass binding assays for live *Y. enterocolitica* were performed in a flow cell. Bacteria were treated as mentioned above, resuspended in microscopy medium at the indicated pH value, and added to the flow cell as described above. Bacteria were tracked visually for 10 min afterwards.

## Acknowledgements

We thank Nicholas Dickenson (Utah State University, Logan, USA) for the kind sharing of *Shigella flexneri* GFP-Spa47, Sophie Bleves (Aix-Marseille Univeristy, FR) for the *Pseudomonas aeruginosa* PAO1 strain, Marc Erhardt (Humboldt University Berlin, Germany), Tohru Minamino (Osaka University, Japan), and Gero Miesenböck (University of Oxford, UK) for the pHluorin_M153R_ DNA, Luke Lavis (Janelia Research Campus, USA) for the kind donation of Janelia Fluor dyes, and Malte Buchholz (University of Marburg, Germany) for helpful discussions about gastrointestinal infection pathways.

This work was supported by the Max Planck Society. It was also funded by the Horizon 2020 Innovative Training Network “ViBrANT” (to DL).

## Footnotes

a In this manuscript, T3SS refers to the virulence-associated T3SS. The common „Sct” nomenclature (Hueck, 1998) is used for T3SS components, see (Diepold & Wagner, 2014) for species-specific names.

b All *Y. enterocolitica* fusion proteins used in this study are expressed from the native genetic environment; the genes replace the wild-type genes by allelic exchange (Kaniga *et al*, 1991).

## Notes

#### Summary of Updates

Additional new results (including new Figure 5 and several Supplemental Figures); results and discussion updated and clarified.

