## Supplementary Information for "Dynamic relocalization of the cytosolic type III secretion system components prevents premature protein secretion at low external pH"

| Strain | Genotype | Reference |
|---|---|---|
| MRS40 | Wild-type pYV *Y. enterocolitica* E40 *ΔblaA* | (Sory *et al*, 1995) |
| IML421*asd* (HOPEMTasd) | MRS40 *yopO*$_{\Delta 12\text{-}427}$ *yopE*$_{21}$ *yopH*$_{\Delta 11\text{-}352}$ *yopM*$_{23}$ *yopP*$_{23}$ *yopT*$_{135}$ *Δasd* | (Kudryashev *et al*, 2013) |
| AD4016 | MRS40 *egfp-sctQ* | (Diepold *et al*, 2010) |
| AD4085 | IML421*asd egfp-sctQ* | (Kudryashev *et al*, 2013) |
| AD4175 | IML421*asd sctV-egfp* (mutated with pAD208) | This work |
| AD4306 | IML421*asd egfp-sctD* | (Diepold *et al*, 2015) |
| AD4411 | IML421*asd egfp-sctQ ΔsctD* | (Diepold *et al*, 2017) |
| AD4439 | IML421*asd pamcherry1-sctD* (mutated with pAD439) | This work |
| AD4474 | IML421*asd efgp-sctK* | (Diepold *et al*, 2017) |
| ADTM4514 | IML421*asd egfp-sctN* | (Diepold *et al*, 2017) |
| ADTM4520 | IML421*asd egfp-sctL* | (Diepold *et al*, 2017) |
| ADTM4521 | IML421*asd mcherry-sctL* | (Diepold *et al*, 2017) |
| ADTM4525 | IML421*asd halo-sctL* | (Diepold *et al*, 2017) |
| ADMH4536 | IML421*asd halo-sctL ΔsctF* | This work |
| DL001 | *P. aeruginosa* PAO1 *egfp-sctQ* | (Lampaki *et al*, 2019) |
| *S. flexneri gfp-sctN* | *S. flexneri* serotype 2a 2457T *gfp-sctN(spa47)* | (Burgess *et al*, 2020) |

| Plasmids | Genotype | Reference |
|---|---|---|
| pBAD-His B | pBR322-derived expression vector | Invitrogen |
| pKNG101 | *oriR6K sacBR+ oriTRK2 strAB+* (suicide vector) | (Kaniga *et al*, 1991) |
| pAD208 | *pKNG101-sctV-egfp* | (Diepold *et al*, 2017) |
| pAD439 | *pKNG101-pamcherry1-sctD* | This work |
| pAD638 | *pBAD::sctF*$_{S5C}$ | This work |
| pEE010 | *pBAD::sctD*$_{H193A,H205S,R214H,H353Y,H376G}$ | This work |
| pISO85 | *pKNG101-ΔsctF* | (Diepold *et al*, 2010) |
| pSW001 | *pBAD::pHluorin* | This work |
| pSW022 | *pBAD::sctD* | This work |

**Supplementary Table 1 – Strains and plasmids used in this study**

| Low salt LB medium | concentration |
|---|---|
| Yeast extract | 5 g/l |
| Trypton | 10 g/l |
| NaCl | 3 g/l |
| Agarose (for plates) | 1.5% |

| BHI medium | concentration |
|---|---|
| Brain heart infusion solids | 17.5 g/l |
| Peptones | 10.0 g/l |
| Glucose | 2.0 g/l |
| Sodium chloride | 5.0 g/l |
| Disodium hydrogen phosphate | 2.5 g/l |
| Agarose (for plates) | 1.5% |

| Minimal microcopy medium | concentration |
|---|---|
| HEPES pH 7.2 | 100 mM |
| $(NH_4)_2SO_4$, ammonium sulfate | 5 mM |
| NaCl | 100 mM |
| Sodium glutamate | 20 mM |
| $MgCl_2$ | 10 mM |
| $K_2SO_4$ | 5 mM |
| MES | 5 mM |
| Glycine | 50mM |
| Casamino acids | 0.5% |
| Agarose (for agarose pads) | 1.5% |

**Supplementary Table 2 – Composition of media used in this study**

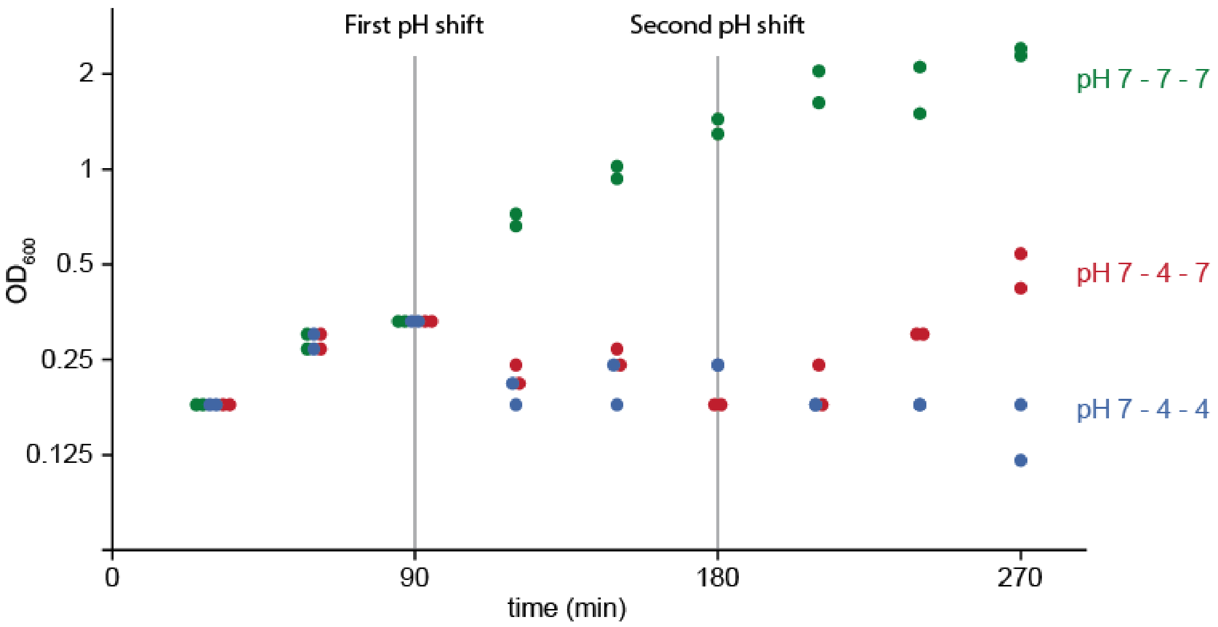


**Suppl. Fig. 1 – *Y. enterocolitica* growth stops at pH 4, and resumes in neutral pH**

*Y. enterocolitica* dHOPEMTasd were inoculated to an $OD_{600}$ of 0.12 from a stationary over night culture. They were grown at pH 7 (28°C) for 90 min (0-90 min), collected by centrifugation and then resuspended in fresh medium (37°C) at pH 7 (green data points) or pH 4 (red and blue data points). Cultures were incubated for another 90 min (90-180 min), again collected by centrifugation, resuspended in fresh medium (37°C) at pH 7 (green and red data points) or pH 4 (blue data points), and incubated for further 90 min (180-270 min). $OD_{600}$ values were recorded every 30 min. The results of two independent experiments are displayed.

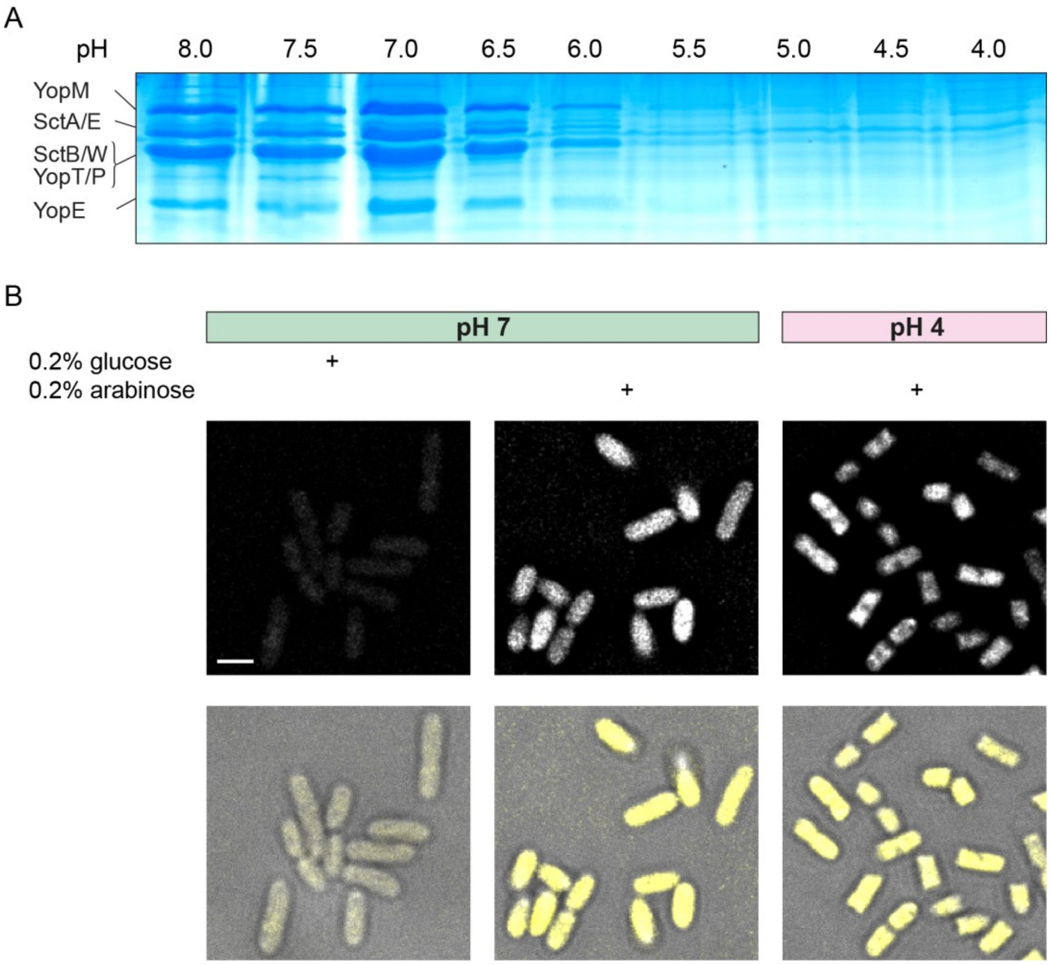


**Suppl. Fig. 2 – Protein export, but not protein synthesis in *Y. enterocolitica* is not suppressed at an external pH of 4**

(**A**) *In vitro* secretion assay showing the export of native T3SS substrates (indicated on left side) in a MRS40-based strain containing all native virulence effectors at the indicated external pH values. Coomassie-stained SDS-PAGE gel; supernatant of $3*10^8$ bacteria per lane. (**B**) *Y. enterocolitica* dHOPEMTasd were grown at neutral pH and then subjected to different pH as indicated. EGFP expression was induced from a pBAD plasmid at the same time, and fluorescence was determined after 180 min. Top, fluorescence image in GFP channel, bottom, overlay of phase contrast (grey) and fluorescence (yellow). Scale bar, 2 µm.

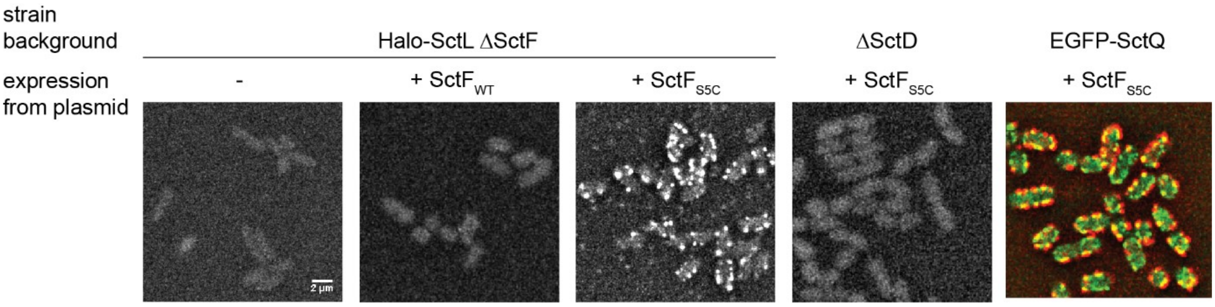


**Suppl. Fig. 3 – Specificity of maleimide-based labeling of SctF$_{S5C}$**

Fluorescence micrographs of *Y. enterocolitica* strains expressing different version of SctF from plasmid, as indicated, induced with 1.0% arabinose. For the four images on the left side, bacteria were stained using CF488A maleimide dye and imaged in the green channel. For the colocalization with EGFP-SctQ (right image), bacteria were stained with CF633 maleimide dye and imaged in the Cy5 and green channels.

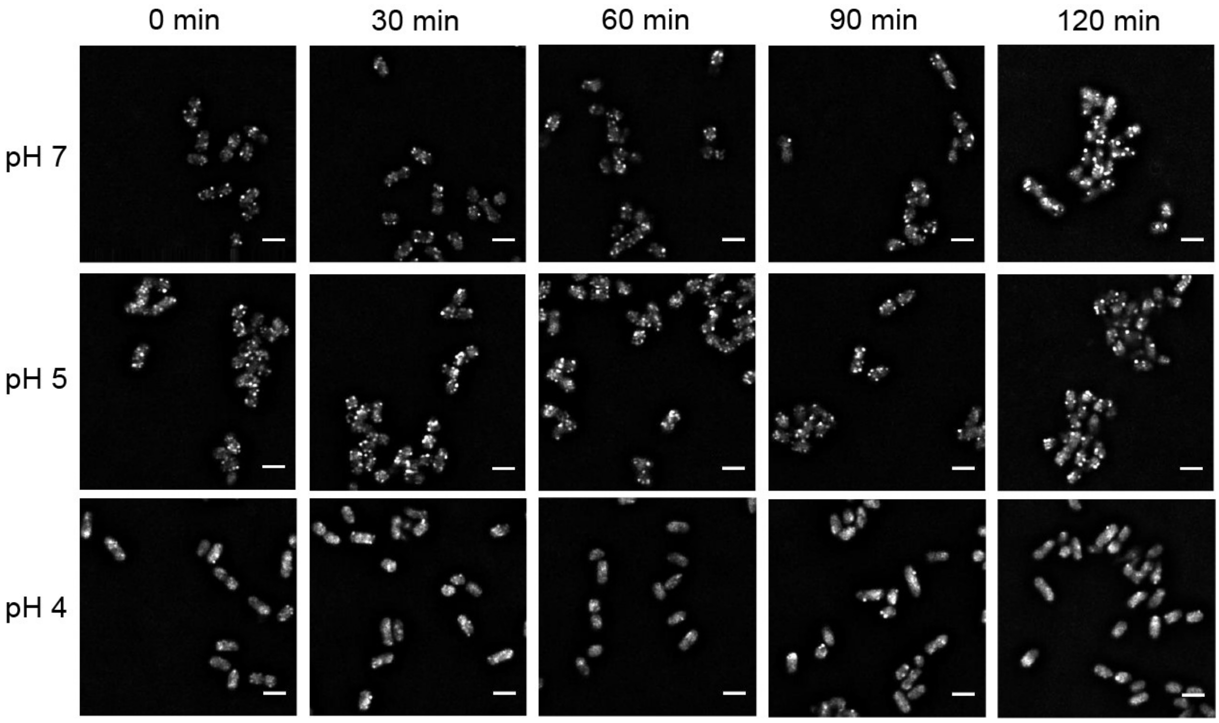


**Suppl. Fig. 4 – The localization of the cytosolic components remains stable over time**

Fluorescence micrographs of *Y. enterocolitica* EGFP-SctQ, incubated at the indicated external pH values under secreting conditions over time. Scale bars, 2 µm.

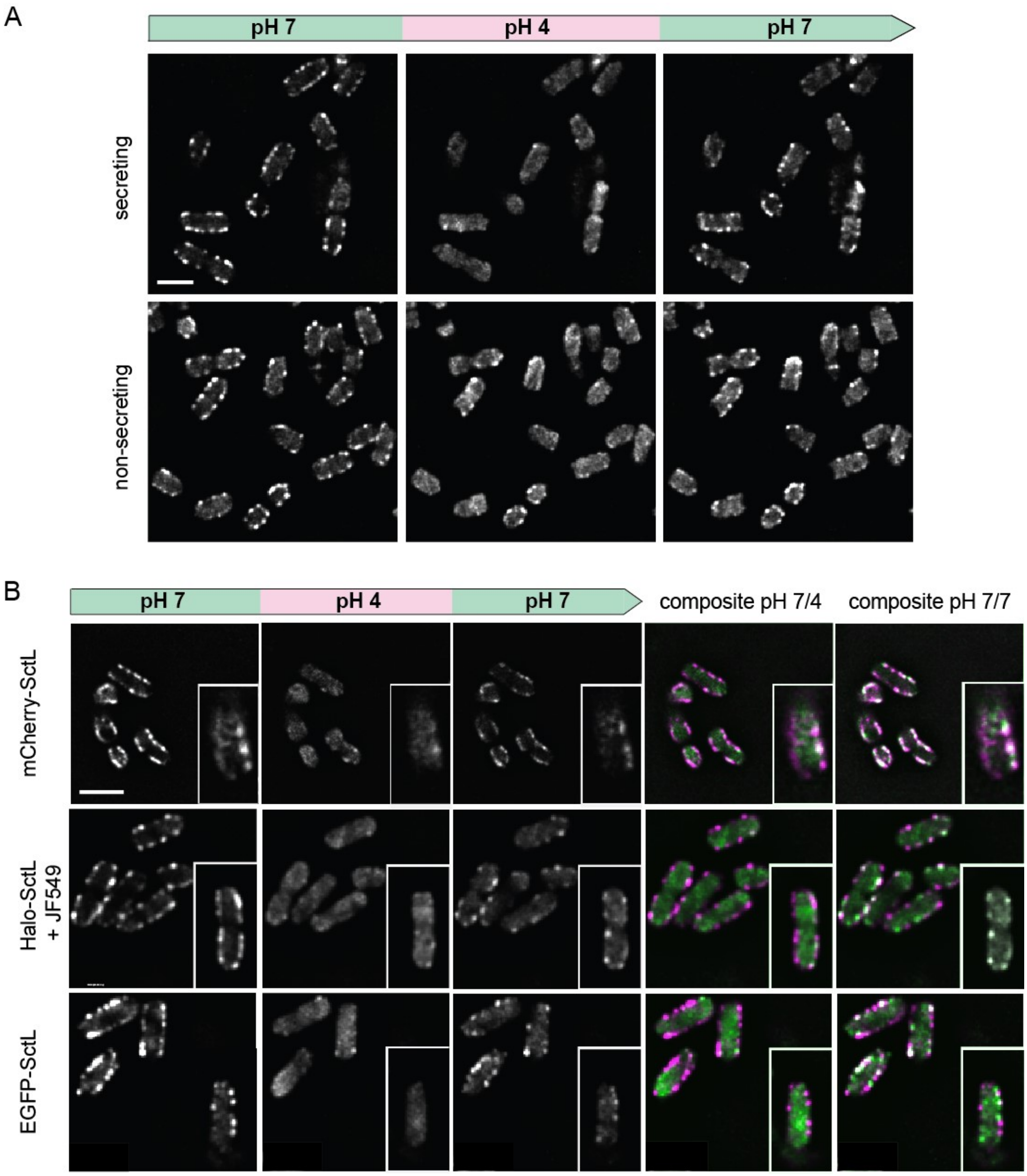


**Suppl. Fig. 5 – Dissociation of the cytosolic components at low external pH can be observed irrespective of the used visualization tag, and in both secreting and non-secreting conditions**

(**A**) Fluorescence micrographs of *Y. enterocolitica* EGFP-SctQ, incubated at the indicated external pH values under secreting conditions (top), or non-secreting conditions (bottom) over time. (**B**) Fluorescence micrographs of *Y. enterocolitica* expressing indicated labeled versions of SctL (replacing the WT gene by allelic exchange) at the indicated external pH values under secreting conditions over time. Right columns, composite images; magenta: pH 7 (first image on the left); green: pH 4 (second image on the left) or pH 7 (third image on the left), as indicated. Scale bars, 2 µm.

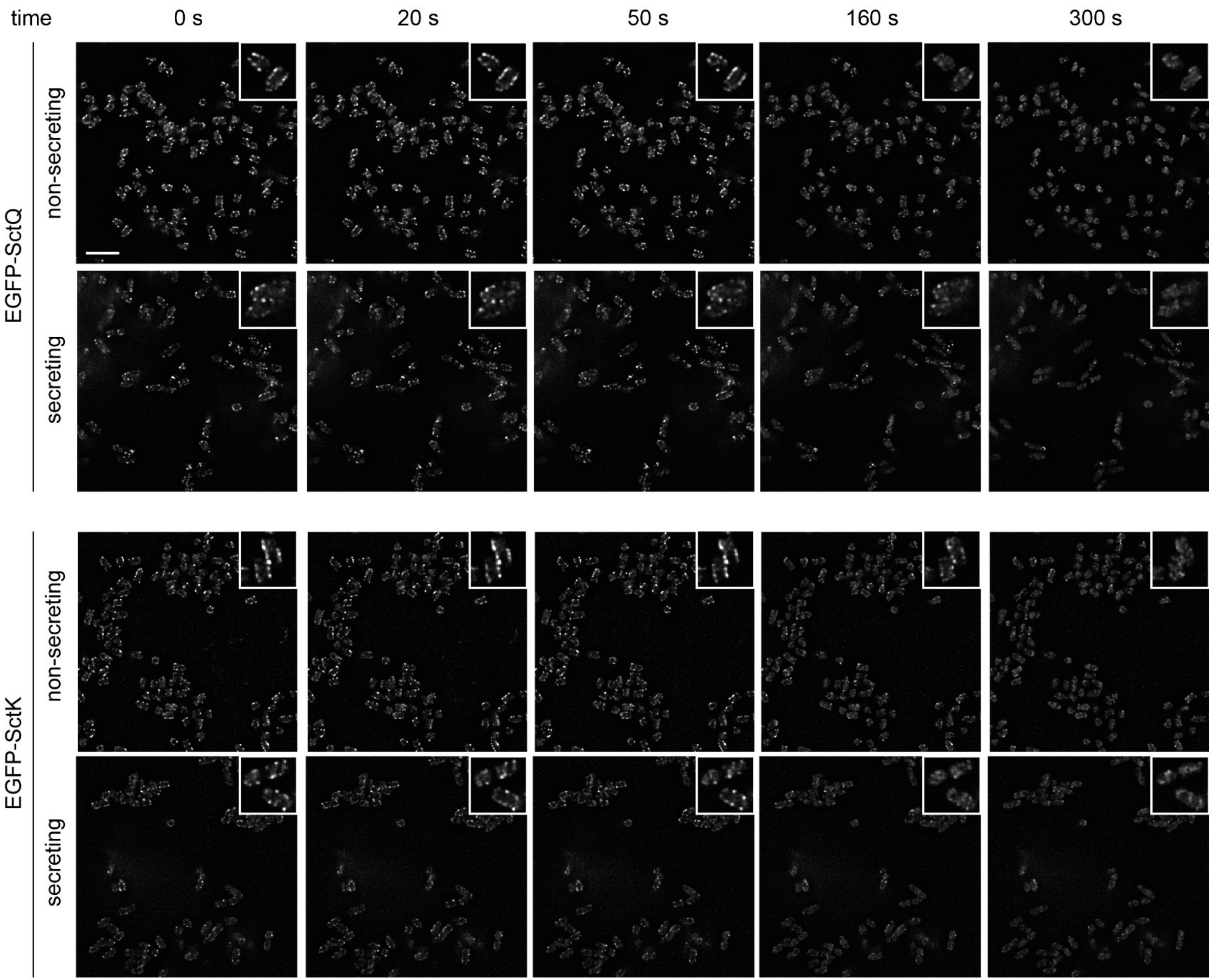


**Suppl. Fig. 6 – Dissociation kinetics of the cytosolic T3SS components under secreting and non-secreting conditions**

Fluorescence micrographs of *Y. enterocolitica* EGFP-SctQ (top) or EGFP-SctK (bottom), at the given time periods after subjecting the bacteria to an external pH of 4 in a flow cell, under secreting conditions (rows 1 and 3), or non-secreting conditions (rows 2 and 4) Scale bar, 5 µm; insets show enlarged sections of the micrographs.

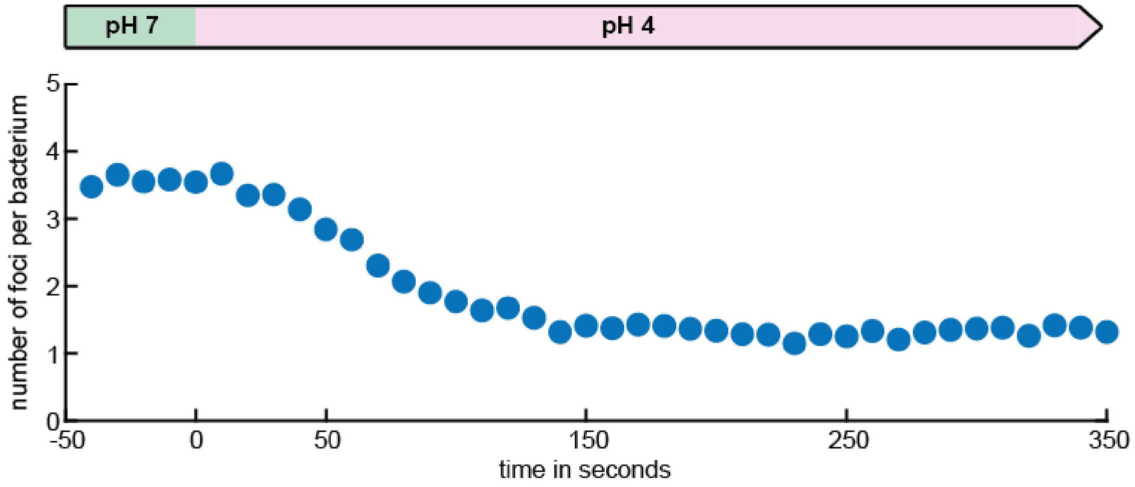


**Suppl. Fig. 7: Quantification of EGFP-SctQ dissociation kinetics upon exposure to external pH 4**

The number of fluorescent EGFP-SctQ foci detected by BiofilmQ (see Material and Methods for details) was determined every 10 s in a flow cell upon changing the external pH from 7 to 4. *n* = 270 bacteria.

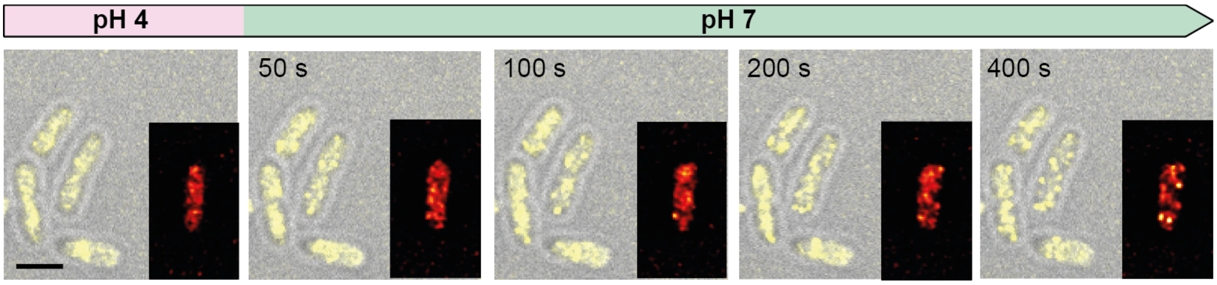


**Suppl. Fig. 8: Re-association kinetics of EGFP-SctQ upon restoration of neutral external pH**

Kinetics of EGFP-SctQ re-association after pH shift from 4 to 7. Overlay of phase contrast (grey) and fluorescence images (yellow); insets, enlarged single bacteria, visualized with the ImageJ red-hot color scale. Scale bar, 2 µm.

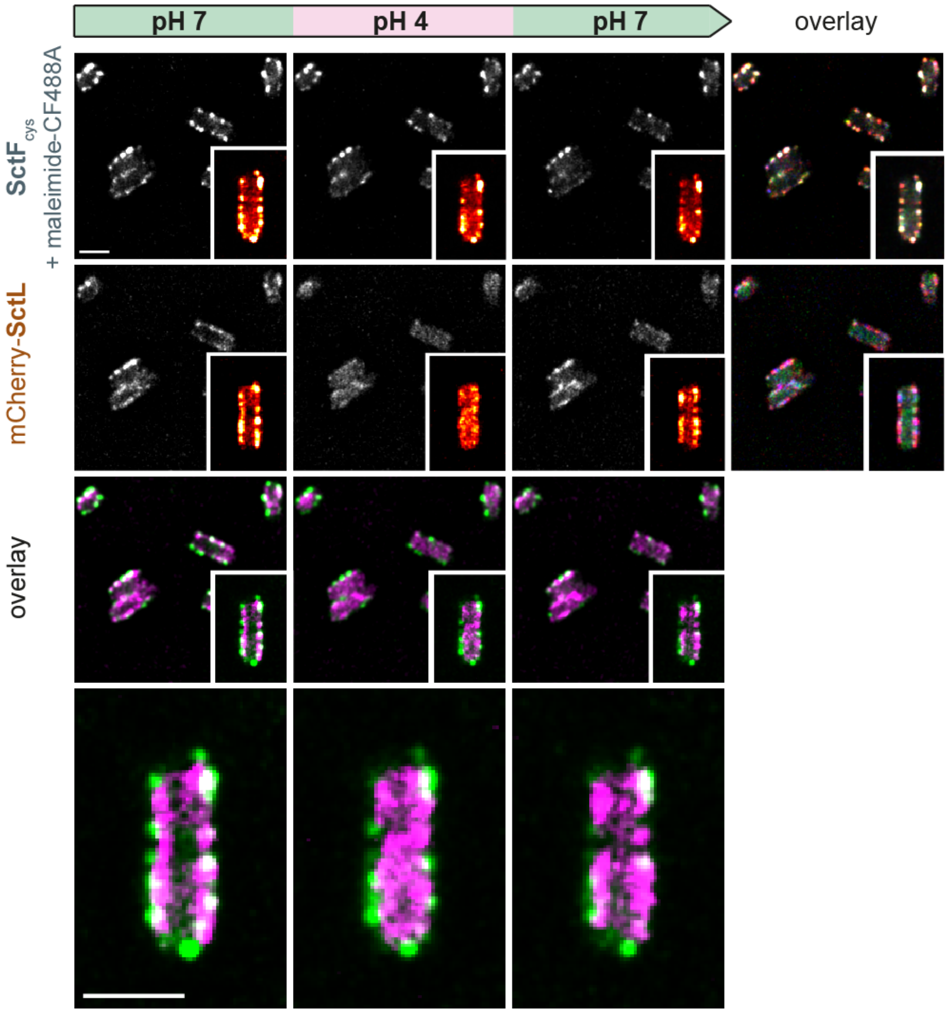


**Suppl. Fig. 9: Cytosolic T3SS subunits colocalize with the needle before and after incubation at pH 4**

Top rows, fluorescence micrographs of *Y. enterocolitica* expressing mCherry-SctL (replacing the WT gene by allelic exchange) and $SctF_{cys}$ (from plasmid) at the indicated external pH values under secreting conditions over time. $SctF_{cys}$ was labeled with maleimide-CF488A and visualized in the green channel. Insets, enlarged single bacteria, visualized with the ImageJ red-hot color scale. Right column, composite images; red channel: pH 7 (first image on the left); green: pH 4 (second image on the left); blue: pH 7 (third image on the left). Third row, composite image of $SctF_{cys}$ and mCherry-SctL (green and magenta, respectively); bottom row, magnification of inset. Scale bars, 2 µm.

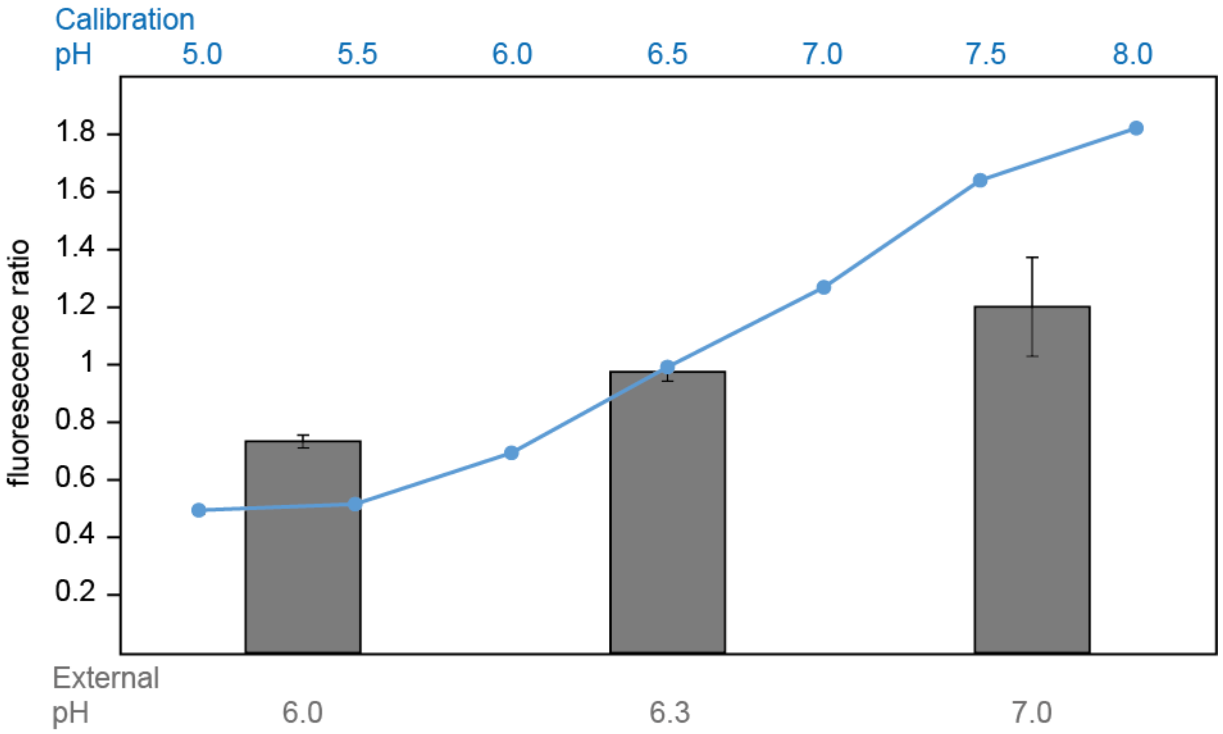


**Suppl. Fig. 10: Internal pH is equilibrated with external pH upon DNP treatment**

Blue curve, calibration of ($Ex_{390nm}$ / $Ex_{475nm}$) fluorescence ratio of purified $pHluorin_{M153R}$ for the pH values indicated on the top (blue) (see Fig. 3A). Technical triplicate, error bars too small to display. Grey bars, determination of cytosolic pH upon incubating bacteria at the indicated external pH values (bottom) in presence of 2 mM DNP. Fluorescence ratio ($Ex_{390nm}$ / $Ex_{475nm}$) of bacteria expressing cytosolic $pHluorin_{M153R}$. *n* = 3, error bars denote standard deviation.

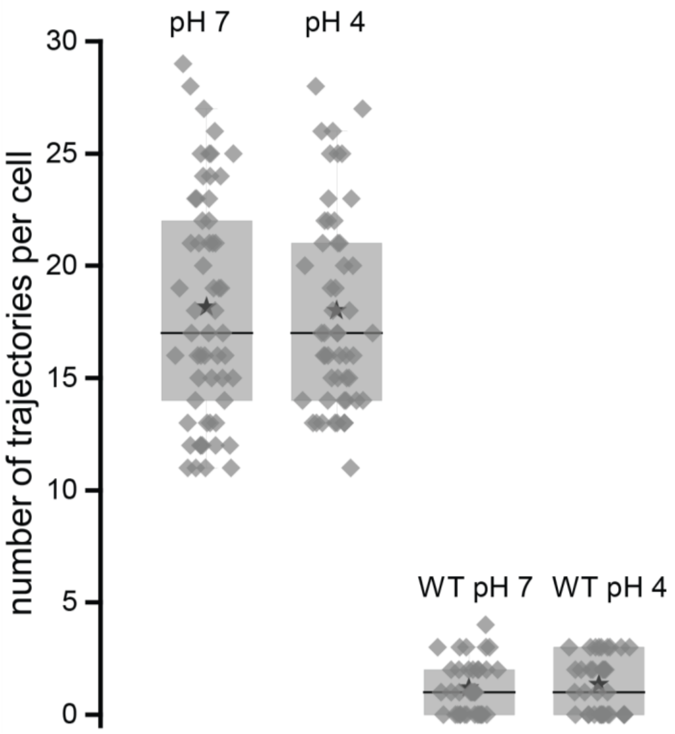


**Suppl. Fig. 11 - Number of SctD trajectories in *Y. enterocolitica* cells at pH 7 and pH 4 compared to the number of false positives measured in wild type cells**

Number of PAmCherry-SctD trajectories per single living *Y. enterocolitica* cells at pH 7 and at pH 4 both exhibit a medium trajectory number of 17 trajectories per cell and a mean of 18.2 ± 5.1 s.d. (pH 7) and 18.0 ± 4.4 s.d. (pH 4). As a control, strains expressing PAmCherry-SctD were mixed with wild type cells during the sample preparation. False positive trajectories from the background signal of single wild type cells in the same movies yield a median of one false positive trajectory per cell for both conditions and a mean of 1.2 ± 1.1 s.d. (pH 7) and 1.3 ± 1.2 s.d. (pH 4). Symbols in the histogram are black star mean, black line median, whisker range 5-95% and box range 25-75%.

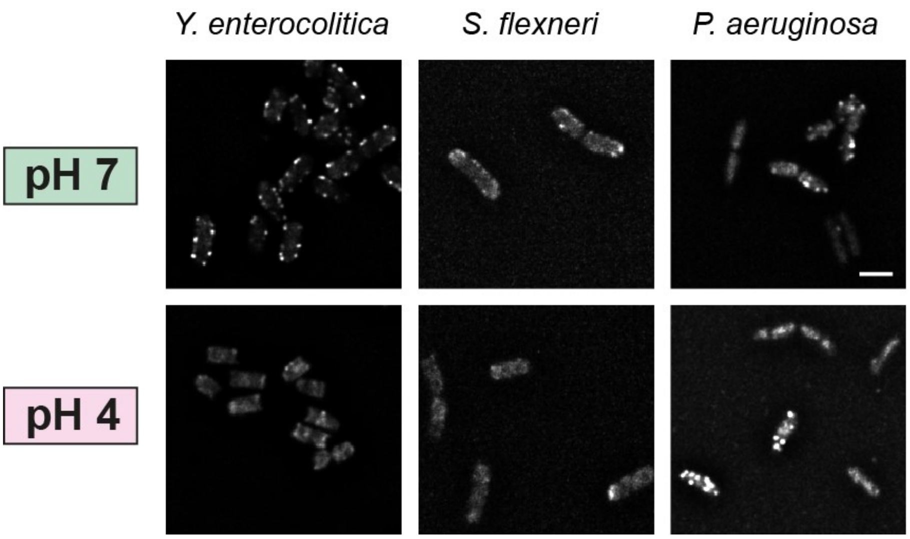


**Suppl. Fig. 12 – Low external pH decreases the fraction of bacteria with foci for fluorescently labeled cytosolic components in *Y. enterocolitica* and *S. flexneri*, but not in *P. aeruginosa***

Representative micrographs of *Yersinia enterocolitica EGFP-SctQ, Shigella flexneri GFP-SctN,* and *Pseudomonas aeruginosa EGFP-SctQ* at the indicated external pH values. Scale bar, 2 µm.

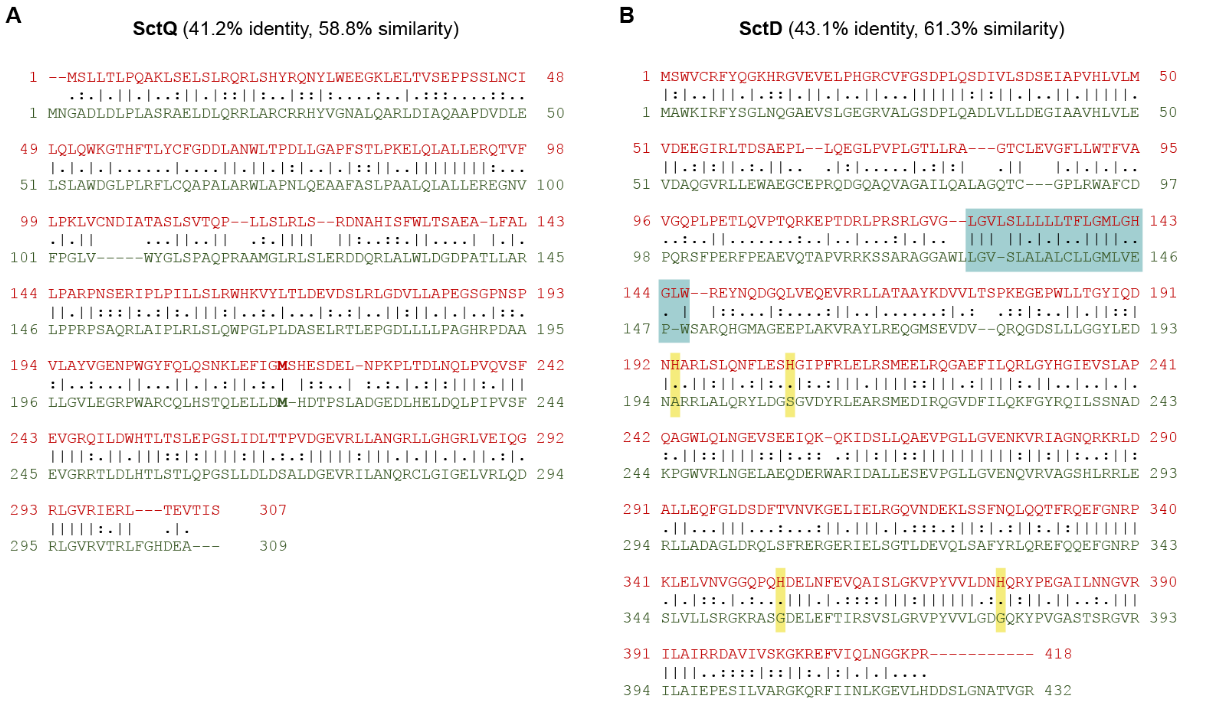


**Suppl. Fig. 13 – Sequence conservation of T3SS components in *Y. enterocolitica* and *P. aeruginosa***

Pairwise sequence alignment of SctQ (**A**) and SctD (**B**). Red, *Y. enterocolitica* sequences (NP_052404, NP_052414); green *P. aeruginosa* PAO1 sequences (NP_250385, NP_250408). Alignments created with EBI EMBOSS Needle (https://www.ebi.ac.uk/Tools/psa/emboss_needle/). Bold M indicates internal translation start site of $SctQ_C$ (amino acid 218 in *Y. enterocolitica*) (Yu *et al*, 2011; Bzymek *et al*, 2012). Turquoise background indicates region of precited trans-membrane helix (TMH) for *Y. enterocolitica* SctD, as predicted by Phobius (Käll *et al*, 2007). The regions upstream and downstream of the TMH correspond to the cytosolic and periplasmic parts of the structure, respectively. Amino acids highlighted in yellow were identified as potentially important for pH sensing in SctD.

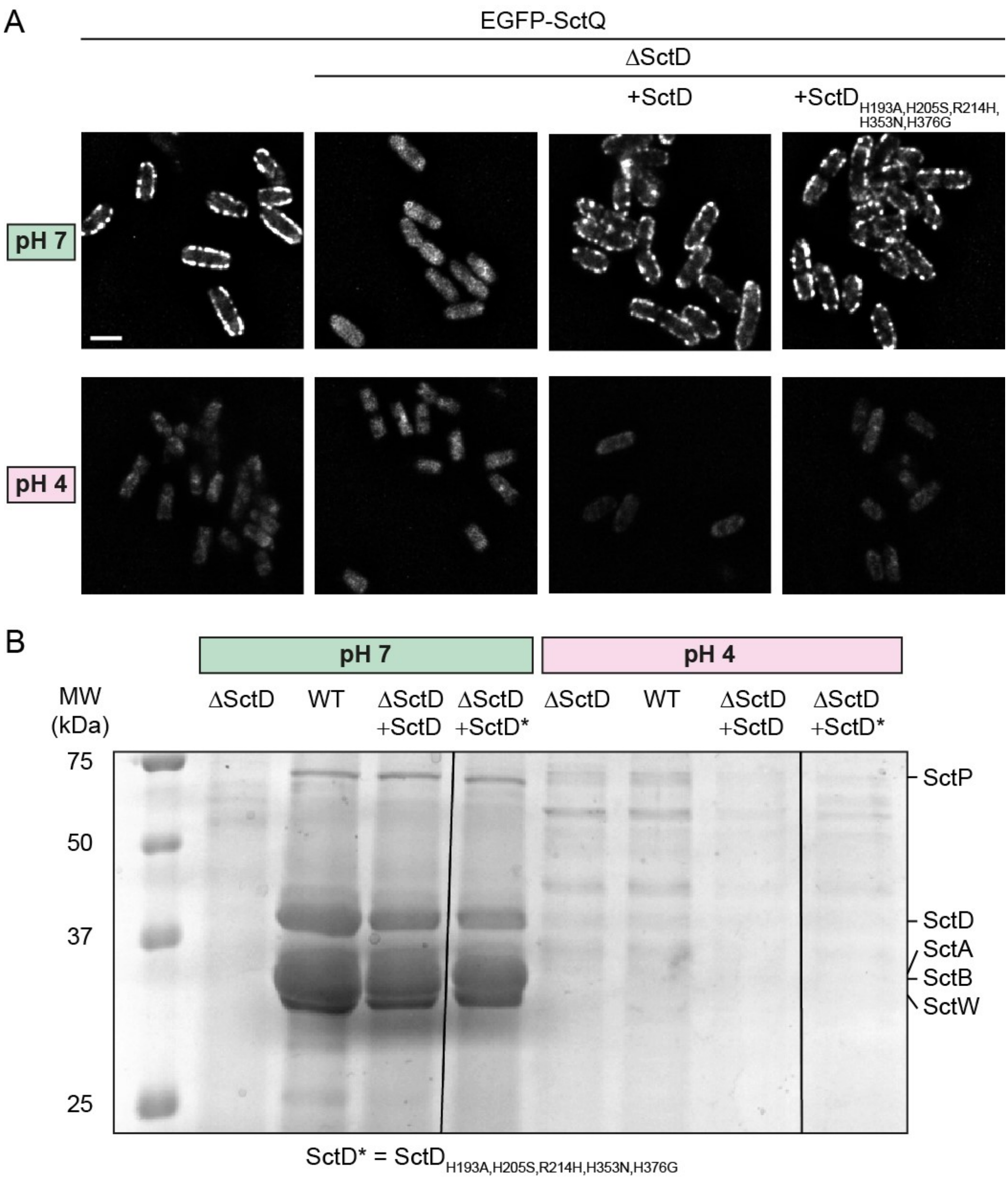


**Suppl. Fig. 14 – Point mutations in SctD do no suppress the pH-dependent dissociation of cytosolic T3SS components and suppression of secretion at low external pH**

(**A**) Fluorescence micrographs of *Y. enterocolitica* EGFP-SctQ in strains lacking SctD (column 2-4) and complemented *in trans* with wild-type SctD (column 3) or an SctD multiple point mutant (see main text for details), at pH 7 (top) or pH 4 (bottom). The mutant SctD confers the same phenotype on SctQ localization as wild-type SctD under both conditions. Sclae bar, 2 µm. (**B**) *In vitro* secretion assay showing the export of native T3SS substrates in the strains used in (A), at external pH of 7 (left) or 4 (right). All samples were analyzed on the same SDS-PAGE gel, vertical lines denote the omission of intermediate lanes. Molecular weight in kDa and exported proteins are indicated and the left and right side, respectively. Black and white scan of a coomassie-stained SDS-PAGE gel; supernatant of $3*10^8$ bacteria per lane.

### Supplementary Videos

#### Suppl. Video 1-2 – *Y. enterocolitica* attaches to surfaces at low external pH

Suppl. Movie 1: Time-lapse phase contrast video of *Y. enterocolitica* attached to a glass cover slip in a flow cell at pH 7. The buffer was exchanged from pH 7 to pH 4 buffered media during the experiment and cells were tracked for 10 minutes with a picture taken every 10 seconds.

Suppl. Movie 2: Time-lapse phase contrast video of *Y. enterocolitica* attached to a glass cover slip in a flow cell at pH 4. During the experiment the buffer was changed from pH 4 to pH 7 and cells were tracked again for 10 minutes with a picture taken every 10 seconds. Scale bars, 2 µm.

#### Suppl. Video 3 – The pH-induced dissociation and re-association of EGFP-SctK to the injectisome can be repeated for several cycles

Time-lapse video of *Y. enterocolitica* expressing EGFP-SctK attached to a glass cover slip in a flow cell. After flow was started the buffer was toggled every 5 minutes between pH 7 to pH 4, as indicated. Micrographs were acquired every 10 seconds. Scale bar, 2 µm.

#### Suppl. Video 4 – Dissociation kinetics of EGFP-SctQ upon change of external pH from 7 to 4

Time-lapse video of *Y. enterocolitica* expressing EGFP-SctQ attached to a glass cover slip in a flow cell after pH shift from 7 to 4. Left, overlay of DIC (grey) and fluorescence signal (yellow); right, fluorescent channel in red hot color scale. Bacteria were attached to the cover slip at pH 7, flow was introduced, and the buffer was switched from pH 7 to pH 4. The duration of the experiment was 10 minutes and pictures were taken every 10 seconds. Scale bar, 2 µm.

#### Suppl. Video 5 – Re-association kinetics of EGFP-SctQ upon change of external pH from 4 to 7

Time-lapse video of *Y. enterocolitica* expressing EGFP-SctQ attached to a glass cover slip in a flow cell after pH shift from 4 to 7. Left, overlay of DIC (grey) and fluorescence signal (yellow); right, fluorescent channel in red hot color scale. Bacteria were attached to the cover slip at pH 7, flow was introduced, and the buffer was switched from pH 4 to pH 7. The duration of the experiment was 10 minutes and pictures were taken every 10 seconds. Scale bar, 2 µm.
